# Rabies virus antagonizes interferon signaling by targeting phosphorylated STAT1 tetramers

**DOI:** 10.64898/2026.07.29.741124

**Authors:** Miku Minami, Aoi Sugiyama, Shatabdi Chakraborty, Tomo Nomai, Cassandra T. David, Yuichiro Takekawa, Fei Yan, Yuta Iizuka, Karyn Wilde, Satomi Inaba-Inoue, Nana Yabuno, Mika Hirose, Tomoyasu Aizawa, Mina Sasaki, Shunsuke Kita, Katsumi Maenaka, Yukihiko Sugita, Gregory W. Moseley, Paul R. Gooley, Toyoyuki Ose

## Abstract

Rabies virus phosphoprotein (P protein) antagonizes interferon signalling by selectively targeting activated STAT1 protein, but the structural basis for the selective recognition of activated STAT1 has remained unclear. Here we report cryo-EM structures of tyrosine-phosphorylated STAT1 (pY-STAT1) in complex with the C-terminal domain of P protein (P-CTD). P-CTD engages the pY-STAT1 tetramer through three discrete interfaces, contacting two DNA-binding domains within a pY-mediated STAT1 dimer and the N-terminal domain of a third STAT1 protomer. This binding mode explains how P-CTD selectively recognizes activated STAT1, occludes surfaces required for importin α5 and DNA binding, and stabilizes the compact tetrameric state, thereby restricting the conformational transition required for cooperative DNA binding. Cell-based assays further indicate that distinct P-CTD surfaces differentially contribute to antagonism of type I and type II interferon signalling. Further, by acting as an anchoring molecule, P-CTD also enabled structural elucidation of the complete STAT1 structures in the tetramer. Together, these data reveal critical mechanisms in viral immune evasion and pathogenesis, and fundamental innate immune signaling.

## INTRODUCTION

The Janus kinase (JAK)–signal transducers and activators of transcription (STAT) pathway mediates cellular responses to more than 50 cytokines via the activity of members of the STAT family of transcription factors, comprising STAT1, STAT2, STAT3, STAT4, STAT5a, STAT5b, and STAT6^1–4^. Many variations of STAT shuttling between the cytoplasm and the nucleus (where STATs activate specific gene expression) have been reported, including trafficking that is dependent or independent of tyrosine phosphorylation, and the formation of distinct complexes including with interferon regulatory factor 9 (IRF9)^5^. Among these mechanisms, the major consensus or paradigm of JAK-STAT signaling is the activation-induced nuclear import of STAT molecules that follows the recognition of specific cytokines by cell surface receptors. Cytokine-bound receptors recruit STATs and receptor-associated JAKs induce phosphorylation of a unique tyrosine residue in the STAT molecules, promoting STAT homo- or heterodimer formation via reciprocal interactions between a phosphotyrosine (pY) and Src homology 2 (SH2) domain in each of the respective STAT molecules^5–7^. Dimerized STATs are then bound by importins, which transport STATs into the nucleus, where they bind to specific promoter regions and regulate the expression of genes involved in cell proliferation, differentiation, and immune responses^3,8,9^. Among the cytokines that activate STATs, the type I IFNs (IFN-α/β) have critical roles in innate immune responses, including antiviral responses; IFNα/β binds and induces heterodimerization of the cell-surface receptors IFN-α/β receptor 1 (IFNAR1) and IFNAR2, leading to the activation of the intracellular receptor-associated JAKs (JAK1) and tyrosine kinase 2 (TYK2)^3,10,11^. This canonically results in the phosphorylation of STAT1 and STAT2, forming pY-STAT1–STAT2 heterodimers which can also associates with IRF9 to form the ternary IFN-stimulated gene factor 3 (ISGF3) complex; within the nucleus ISGF3 binds to interferon-stimulated response elements (ISREs) in the promoters of IFN-regulated genes (ISGs) to control their expression^10,12,13^. In contrast, in the type II IFN pathway, IFN-γ is recognized by interferon gamma receptor 1 (IFNGR1) and IFN–γ receptor 2 (IFNGR2), leading to the activation of JAK1 and JAK2^14^. These JAKs phosphorylate STAT1, leading to the formation of a pY-STAT1 homodimer^15^ (and N-terminal domain (NTD)-tethered pY-STAT1 tetramer^16^, below). pY-STAT1 homodimer, known as gamma interferon-activated factor (GAF), translocates to the nucleus to bind to gamma interferon-activated sequences (GAS) within the promoters of IFN-γ-responsive genes^17,18^. ISRE and GAS elements cooperatively regulate the expression of hundreds of IFN-stimulated genes (ISGs) and contribute to antiviral responses^19^. Type I IFN signaling can also induce homo-dimerization of pY-STAT1 to activate GAF–GAS signaling in addition to the major response *via* ISGF3–ISRE signaling^20,21^.

The biology and structural details of classical pY-STAT1 homodimers are well researched, but activated STATs also undergo further regulation, including via STAT oligomerization beyond simple dimerization. STAT tetramerization can have significant impact on DNA binding, referred to as cooperative DNA binding, for example through binding to sequences arranged in tandem. This oligomerization has been reported for various STAT members, but is most commonly associated with pY-STAT1, *via* interactions of the N-terminal domain of pY-STAT1 dimers^22–25^. We reported the first tetrameric structure of pY-STAT1 and proposed a transition model for tandem GAS recognition ^16^. In this model, two STAT1 dimers simultaneously engage distinct GAS motifs on promoter DNA, enabling cooperative regulation of antiviral gene expression. This tetrameric assembly provided new insights into the molecular mechanisms underlying STAT-mediated transcriptional specificity and suggests additional layers of control in IFN responses. In our previous report, however, we could not locate the NTD of STAT1 in the cryo-EM map because of the intrinsically disordered loop connecting the NTD and the core region (the core region comprises a coiled-coil domain (CCD), a DNA-binding domain (DBD), a linker domain (LD), a SH2 domain, and a part of the transactivation domain (TAD)) (Fig. 1a). Therefore, different experimental approaches including the analysis of alternative molecular complexes, are needed to capture the complete tetrameric structure of STAT1 molecules and elucidate the biophysical relevance of STAT tetramerization in cells.

**Fig. 1.**
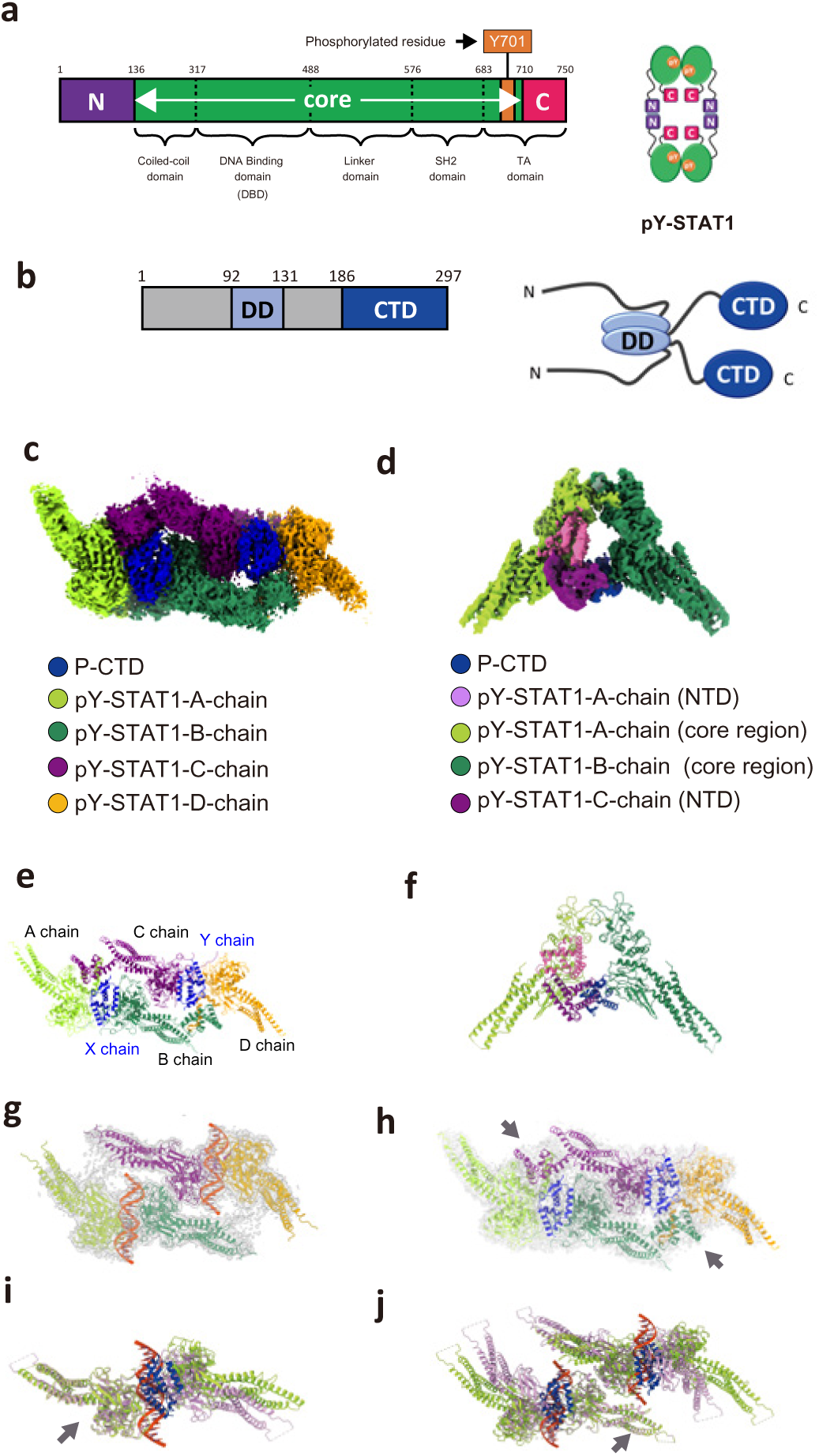
Domain structures of pY-STAT1 and P-CTD, and cryo-EM structure of the pY-STAT1–P-CTD complex. **(a)** Schematic of the domain structure of human STAT1 and the oligomerization states of pY-STAT1. Conserved domains and Tyr701 (Y701, the phosphorylation site) are indicated. The core region comprises the coiled-coil, DNA-binding, linker, and SH2 domains, together with a portion of the TA (transactivation) domain. N indicates the N-terminal domain and C indicates the C-terminal region. pY-STAT1 forms a tetramer via interactions of the pY and SH2 domains, and of the N-terminal domains. **(b)** Schematic of the domain structure of RABV P protein and its oligomerization states. DD indicates the dimerization domain and CTD indicates the C-terminal domain. P protein forms a dimer via the DD interactions shown. **(c)** EM density (3.7 Å resolution) of tetrameric pY-STAT1 in complex with P-CTD. The density corresponding to the A–D chains of STAT1-core region are colored in light green, green, purple, and orange respectively. The P-CTD is colored in blue. **(d)** EM density (3.1 Å resolution) focusing on pY-STAT1-core dimer composed of the core regions of the A (light green) and B (green) chains. The NTD dimer composed of the A (pink) and C (purple) chains, and P-CTD (blue) are also shown. **(e)** Ribbon diagram of the tetrameric pY-STAT1 in complex with P-CTD representing the assembly of each protomer. The A–D chains of STAT1 (NTD–core) are colored in light green, green, cyan, and orange respectively. Two P-CTDs are named X and Y chains (both colored in blue). **(f)** Ribbon diagram of pY-STAT1-core dimer composed of A (light green) and B (green) chains of which the EM density is shown in **d**. NTD dimer composed of A (pink) and C (purple) chains and P-CTD (blue) are also shown. **(g)** EM density and ribbon diagram of tetrameric pY-STAT1 (A–D chains, colored as in **c**) in complex with DNA (previous study^16^, DNA is colored red). **(h)** EM density and ribbon diagram of tetrameric pY-STAT1 (A–D chains, colored as in **c**) in complex with P-CTD (this study). The gray arrows indicate NTD dimers. **(i)** Comparison of pY-STAT1-core dimers between pY-STAT1 (purple) in complex with DNA (red), and pY-STAT1 (green) in complex with P-CTD (blue). The superposition was carried out based on the protomer located on the left (indicated with a gray arrow). **(j)** Comparison of pY-STAT1-core tetramers between pY-STAT1 (purple) in complex with DNA (red) and pY-STAT1 (green) in complex with P-CTD (blue). The superposition was carried out based on the protomer indicated with a gray arrow.

Due to the antiviral roles of IFN-activated JAK-STAT signaling, antagonism of this system is critical in infection by diverse viruses, enabling subversion of host cell biology and immunity. Many viruses make proteins called IFN antagonists, including structural or non-structural viral proteins that interact with and inhibit STAT proteins, representing a critical aspect of host-pathogen dynamics and pathogenesis, and one of the key determinants of viral tropism for cell types, tissues, and species^26^. The virus–STAT interface is thus considered to be a potential target for the development of antivirals and live-attenuated safe vaccines^27^. Due to the ability of viral proteins to specifically interact with and manipulate STAT functions, characterization of viral IFN antagonist-STAT complexes can also provide probes to understand STAT biology^27^. Thus, structural determination of virus–STAT interactions can inform on viral pathogenesis, interventions, and fundamental processes in cell biology; however, there are only a limited number of recent structural studies of viral proteins with STAT1 or STAT2, specifically on complexes of the C protein of Sendai virus (SeV-C), the 018 peptide of Vaccinia virus, and the NS2A and NS5 proteins of Zika and Dengue viruses, and these studies have used fragments of the STAT, and have not assessed full STAT complexes, thus providing potentially incomplete data^28–32^.

Rabies virus (RABV), a causative agent of rabies disease, is a member of the order *Mononegavirales*, which comprises viruses with a non-segmented, negative-sense single-stranded RNA genome. RABV belongs to the genus Lyssavirus and is responsible for severe zoonotic neurological disease with a case fatality rate approaching 100%, accounting for approximately 59,000 reported human deaths annually across over 150 countries^33^. The RABV genome encodes five structural proteins: nucleoprotein (N), phosphoprotein (P), matrix protein (M), glycoprotein (G), and the large (L) RNA-dependent RNA polymerase^34^. Studies using chimeric virus strains have demonstrated that the N, M, and P proteins are key contributors to RABV pathogenicity^35^. The P protein of RABV (RABV-P) is highly multifunctional, with roles including acting as the essential cofactor of the viral polymerase L protein, and as a potent antagonist of host innate immune responses^36,37^, including through interaction with and inhibition of the STAT proteins, including STAT1, 2, and 3^38–41^.

Several cell-based assays support roles for RABV-P in immune evasion, suggesting that it interacts with and inhibits signaling by phosphorylation-induced pY-STAT1 homodimer/homotetramer, pY-STAT1–STAT2, and pY-STAT1–STAT3 heterodimers, but not the pY-STAT3 homodimer^38,39,41^. RABV-P is also expressed as multiple N-terminal truncated isoforms that all share a common dimerization domain (residues 92–131) and C-terminal domain (P-CTD; residues 186–297) (Fig.1b)^42^. For the inhibition of type I and type II IFN signaling, truncation studies have indicated that the P-CTD directly interacts with STAT1^36^. This inhibition involves the cytoplasmic sequestration of STAT1 as well as disruption of the STAT1–DNA interaction^36,41^. Notably, these mechanisms of STAT1 inhibition are conserved among lethal lyssavirus species^42^. Mutagenetic analysis of P-CTD initially suggested that residues Trp265 and Met287, located within a hydrophobic region and called “W-hole”, are involved in STAT1 or STAT2 interaction and the suppression of the type I IFN response^43^. Comparative structural studies suggested that the bulkiness of the hydrophobic surface at position 265 correlates with viral pathogenicity^44^. Extensive molecular dissection by NMR spectroscopy, however, identified three distinct regions of P-CTD (none of which included Trp265 or Met287) as candidates for the direct interaction with STAT1^45^, suggesting that the effects of Trp256 and Met287 mutation may be indirect, affecting the stability of the P-CTD. Among the residues in the regions identified by NMR, a mutation-scanning–based screen using a reporter gene assay demonstrated that a single mutation of Phe209, Asp235, as well as Trp265, or Met287 generated moderately defective P protein for inhibition of IFN-α induced STAT signalling^45^. It has been suggested that P-CTD can recognize the phosphorylation state of STAT1 and primarily targets pY-STAT1 in host cells, as binding of P protein to STAT1 detected by approaches such as immunoprecipitation is dependent on activation by type I or type II IFN^36,38,43,45^. Indeed, quantification of dissociation constants (*K*_D_ values) using purified proteins revealed that P-CTD binds to pY-STAT1 much more strongly than to U-STAT1, indicating a viral strategy to selectively target activated STATs, thereby disabling innate immune functions, but not affecting other potential functions of U-STATs^16^. P-CTD additionally recognizes the presence of the NTD of pY-STAT1^16^. P-CTD is also capable of directly interacting with the nucleoprotein (N protein) and appears to occur using a site that appears distinct from the STAT1 binding sites^35^. Based on these findings, we previously predicted a binding mode of the pY-STAT1–P-CTD complex^16^. As mutations affecting interactions between STAT molecules and P have been shown to reduce viral pathogenesis, detailed information on the pY-STAT1–P-CTD interaction could provide insights contributing to the development of attenuated vaccine strains, or therapeutics to disrupt the interaction^45^; however, our prediction model requires direct experimental assessment before such advances are possible.

Here we report the cryo-EM structure of tetrameric pY-STAT1 in complex with P-CTD. This structure reveals how the P protein selectively targets pY-STAT1 in IFN-activated cells by recognizing the pY–SH2 interaction-mediated dimeric assemblies: specifically, P-CTD interacts with two DBDs of two pY-STAT1 core regions related with a two-fold symmetry, which will inhibit the association of STAT1 with importin α5 (which mediates nuclear import of the activated pY-STAT1 complexes) and with promoter DNA. Furthermore, the P-CTD interacts with the N-terminal domains (NTD) of the third STAT1 protomer; thus, P-CTD recognizes tetramerization as well as the dimerization of pY-STAT1. Together, these data indicate that the P-CTD engages pY-STAT1 through three distinct interaction sites that are spatially and functionally separate from the P-CTD–N-protein binding site. Quantitative analysis of dimeric and tetrameric pY-STAT1 further indicates that P-CTD stabilizes the tetrameric state and suppresses the conformational transition required for cooperative DNA binding, revealing a previously unrecognized mechanism of STAT antagonism. Notably, by combining these structural insights with reporter gene assays comparing the effects of P-CTD mutations on the type I (STAT1–2 dependent) and II (STAT1) IFN pathways, this structural framework offers a mechanistic explanation for how P-CTD can also recognize STAT1-containing heterodimers, including STAT1–STAT2 and STAT1–STAT3^36,39,40^. These data provide major insights into the immune evasion and pathogenic mechanisms of RABV. In addition, because two P-CTDs bind a tetramer of pY-STAT1 and act as molecular anchors that immobilize the NTDs of the pY-STAT1 tetramer, its visualization became enabled in the cryo-EM structure for the first time. These data directly confirm that two NTDs tether two pY-STAT1 dimers to form a compact tetrameric architecture, providing new insights into the nature of the physiological STAT1 tetrameric structure. As the intact form of P protein forms a dimer, the 4:2 complex formed by P-CTD binding means a single P dimer binds to the single functional STAT hetero- or homotetramer.

## RESULTS

### Overall structure reveals one tetrameric pY-STAT1 is recognized by two P-CTD molecules

We previously reported the tetrameric pY-STAT1 structure (*C*2 of *C*2 symmetry) in complex with DNA using cryo-EM^16^. Herein, we reconstructed 3D cryo-EM maps of homotetramers and homodimers of pY-STAT1 in complex with the P-CTD of the Nishigahara (Nish) RABV strain P-CTD (Fig. 1c,d, Extended Data Fig.1, 2, 3A, B). The resolution of the final map was determined using the Fourier shell correlation (FSC) method at the FSC 0.143 criterion, achieving a resolution of 3.7 and 3.1 Å respectively. Subsequently, the atomic model of the pY-STAT1 tetramer in complex with the P-CTD was refined against the final map (Fig. 1c, e). In the EM density reconstructed for the dimeric pY-STAT1 core regions (A and B chains), clear density also covering P-CTD (X chain) and two NTDs of pY-STAT1 (an NTD dimer composed with the A and C chains, see below) were observed (Fig. 1d, f); each pY-STAT1-core is stabilized by the reciprocal pY-SH2 interactions as observed in the dimeric pY-STAT1 structures in complex with DNA^6,7,16^.

In the density for tetrameric pY-STAT1, four protomers of the pY-STAT1-core were easily located based on the analysis using the model built in the dimeric form: there are two pairs of pY-STAT1-core dimers (AB dimer and CD dimer (Fig. 1c)). These two dimers assemble into a tetramer through two pairs of NTD–NTD interactions: between A–C protomers and between B–D protomers (Fig. 1c, Extended Data Fig.3C). The map quality of the D chain was the least well resolved; the D chain has the greatest number of truncated residues in the map. Notably, while the NTDs could not be located in the previous pY-STAT1–DNA structure (Fig. 1g), we observed density corresponding to the NTDs in the pY-STAT1–P-CTD map in the current study (Fig. 1h).

As previously observed in the cryo-EM structure of pY-STAT1–DNA^16^, for the tetrameric pY-STAT1 structure bound to P-CTD, the density revealed the phosphorylated tyrosine (Tyr701) and the recognition of this phospho-moiety by the SH2 domain of the opposing protomer using residues around Lys584 and Arg602 (between A–B and C–D protomers in in each of the dimers, Extended Data Fig.3D). However, the overlay on the A protomers of the two structures (pY-STAT1–DNA and pY-STAT1–P-CTD structures) indicates that the relative angle for the opposing B protomer is different: i.e. the two protomers of STAT1-core are not related by pure two-fold rotation symmetry in the pY-STAT1–P-CTD dimer structure in contrast to the pure two-fold rotational symmetry in the pY-STAT1–DNA structure (Fig.1i). By contrast, the STAT1-core regions of the B and C protomers are related with pseudo two-fold rotation symmetry in the pY-STAT1–P-CTD structure, consistent with that observed in the tetrameric structure of pY-STAT1–DNA^16^, although *C*2 symmetry averaging was not applied for the map refinement. The NTD of the C-chain makes contact with the DBD of the A chain and P-CTD (X chain) and dimerizes with the NTD of the A-chain, enabling the visualization of the NTD dimer ofpY-STAT1 (A and C chains, Fig. 1c, h). There is no density for the C-terminal region of STAT1 (all chains, residues 720–750).

Density corresponding to the two P-CTD molecules (X and Y chain) was observed between two DBDs of the reciprocal pY-SH2 tethered dimers (AB and CD dimers of STAT1-core) (Fig.1c), indicating a role as a bridge between two protomers in each pY-STAT1 dimer (Fig. 1c, e). Structural comparison of the pY-STAT1-core dimers and three-dimensional variability analysis (3DVA) revealed that there is a hinge like movement between pY-STAT1-cores^16^. This mobility is utilized for the induced-fit P-CTD binding to the pY-STAT1-core dimer. This demonstrates a binding stoichiometry of 4:2 (pY-STAT1:P-CTD) as predicted previously^16^. As P protein forms a dimer using the DD (Fig. 1b), the stoichiometry proposes that one dimer of P protein targets one tetramer of pY-STAT1; in common with the P-CTD, the DD is conserved in all isoforms of P protein, so similar stoichiometry would be expected. Using small angle X-ray scattering (SAXS) experiments, we previously reported that large conformational changes do not happen in solution among the pY-STAT1 tetramer (the free form), the pY-STAT1 tetramer in complex with two double-stranded 18mer DNA molecules, and the pY-STAT1 tetramer in complex with P-CTD^16^. Additionally, using the cryo-EM structure of pY-STAT1–DNA, we previously proposed a transition model of tetrameric pY-STAT1 from the compact packed form (e.g., the free form) to the tandemly promoter-bound form^16^. Besides the hinge-like movement between the cores, the overall conformation of tetrameric pY-STAT–P-CTD structure is actually similar to that of tetrameric pY-STAT1–DNA (the compact packed form) as predicted from the SAXS experiment and binding assays^16^. Thus the cryo-EM structure of pY-STAT1–P-CTD indicates that this structure transition is inhibited by the binding of the two P-CTDs to the compact-tetrameric form with an extensive surface of contact (see below), which proposes a novel mechanism whereby P protein prevents STAT-DNA interaction to antagonize signaling.

### pY-STAT1 and P-CTD interact via three different binding interfaces

The position of the P-CTD in the complex reveals three discrete binding interfaces, designated sites A, B, and C according to the STAT1 protomer chains engaged at each site. Each site on the P-CTD interacts with a different STAT1 protomer: site A with the core of the A protomer of the pY-STAT1 tetramer (pY-STAT1-A-core), site B with the core of the B protomer (pY-STAT1-B-core), and site C with the NTD of the C protomer (pY-STAT1-C-NTD) (Fig. 2a). The fact that P (specifically P-CTD) extensively binds to pY-STAT1 in the context of phosphorylated multimers agrees well with the previous observation that efficient binding between P-CTD and STAT1 is promoted after IFN stimulation^36,38,43,45^. Previously using NMR and reporter gene assay following structure-guided mutagenesis, we reported that three distinct regions on the P-CTD surface are important for STAT1 binding; two of which interact with STAT1-CCD-DBD, whereas the other interacts with the STAT1-NTD, or the STAT1-C-terminal region^45^. We also suggested that P-CTD recognizes pY-SH2 mediated dimerization as well as NTD-tethered tetramerization and built a 4:2 binding model of pY-STAT1 and P-CTD^16^. Thus, the experimental cryo-EM structure of pY-STAT1 tetramer in complex with P-CTD generated here is consistent with the accumulated prior data regarding the interaction.

**Fig. 2.**
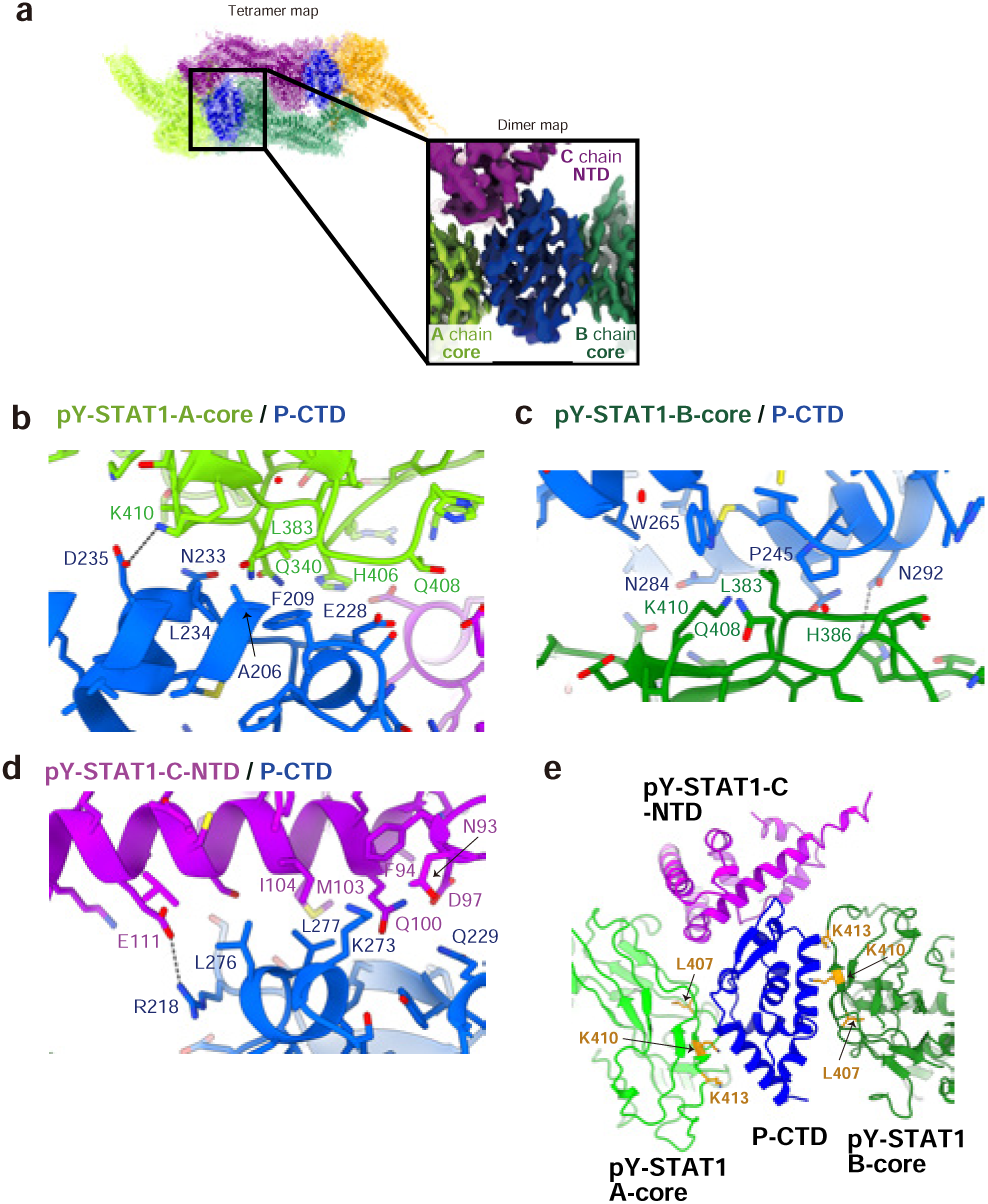
Molecular basis of the interaction between pY-STAT1 and P-CTD. **(a)** EM map and ribbon diagram of the tetrameric pY-STAT1 (A–D chains, colored as in Fig. 1c, in complex with P-CTD (blue); inset expands the map and model, highlighting the contacts between the pY-STAT1 chains and domains with the P-CTD. **(b)-(e)** Close-up view showing interaction between pY-STAT1 and P-CTD. Salt bridges and hydrogen bonds are shown as dashed lines. **(b)** pY-STAT1-A-chain-Core/P-CTD interaction. **(c)** pY-STAT1-B-chain-Core/P-CTD interaction. **(d)** pY-STAT1-C-chain-NTD/P-CTD interaction. **(e)** P-CTD interaction with the importin binding site of STAT1 (A and B chains). STAT1 residues (L407, K410, and K413), reported as a dimer-specific NLS, are highlighted in orange. P-CTD is in blue, pY-STAT1 A chain core in light green, B chain core in green, and C chain NTD in magenta.

### The dimerization of pY-STAT1 is recognized by the P-CTD sites A and B

At the pY-STAT1-A-core/P-CTD interface, the region spanning from helix α1 to β-sheet β1 and the region from helix α2 to α3 of the P-CTD interact with a pY-STAT1-DBD. This interaction involves residues Ala206, Phe209, Glu228, Lys231, Asn233, Leu234, and Asp235 of P-CTD, and residues Gly338, Gln340, Leu383, Gly384, Thr385, His406, Gln408, and Lys410 of pY-STAT1-DBD of protomer A (Fig. 2a, b, Extended Data Fig. 4A–C). Regions previously implicated by NMR and mutagenesis as important for STAT1 targeting by the P-CTD, and considered as distinct two regions around Phe209 and Asp235^45^, are now proved to be included in the binding site A. Residues Ala206 and Phe209 of the P-CTD provide the hydrophobic interactions with residue Leu383 in the DBD of pY-STAT1 (Extended Data Fig.4A). In parallel, the carboxylate moiety of Glu228 of the P-CTD contributes the π–π interaction with the peptide bond between Gly384–Thr385 of pY-STAT1 (Extended Data Fig.4A). The negatively charged side chain of Asp235 of the P-CTD forms salt bridges with the positively charged side chain of Lys410 of pY-STAT1 (Extended Data Fig.4B). Additionally, the backbone carbonyl oxygen atom of Met232 of the P-CTD forms a hydrogen bond with the carbamoyl moiety of Gln408 of pY-STAT1 (Extended Data Fig.4C). The main chain amide group of L234 forms a hydrogen bond with the main chain carbonyl oxygen atom of Gly338 (Extended Data Fig.4C). The side chain of Lys231 interacts with the side chain of His406 (Extended Data Fig.4C). The significance of Phe209 and Asp235 of P-CTD for the interaction with STAT1 and/or STAT2 has been previously proposed^45^. The importance of this site is supported by the double mutations of Phe209 and Asp235 which abolished the ability of P protein to inhibit signaling in an IFN-STAT1/2-dependent reporter gene assay, and resulted in attenuation of a recombinant rabies virus^45^.

The regions spanning from helix α3 to α4, as well as helices α5 and α6 of the P-CTD form the pY-STAT1-B-core/P-CTD interface. This interaction involves residues Pro245, Gly246, Trp265, Asn284, Met287, Gln288, and Asn292 of P-CTD, and residues Gln340, Leu383, Gly384, Thr385, His386, His406, Gln408, Lys410, and Gln412 of pY-STAT1-DBD of protomer B (Fig. 2c, Extended Data Fig.4D–F). A side-chain specific hydrogen bond is formed between Asn292 of the P-CTD and His386 of pY-STAT1-B (Extended Data Fig.4D). The carbamoyl moiety of Q288 forms a hydrogen bond with the main chain carbonyl oxygen atom of Leu383 (Extended Data Fig.4D).The hydrophobic patch around W-hole formed by Trp265 and Met287 of the P-CTD^43,46^ appears to engage in hydrophobic interactions with Leu383 and Lys410 (the alkyl chain) of the pY-STAT1 B protomer (Fig. 2c, Extended Data Fig.4E). The importance of the bulkiness of the sidechain corresponding to Trp265 and Met287 in the hydrophobic patch of P-CTD has been studied^44,45^. Recombinant RABV with the W265G/M287V mutations generated attenuated virus^43^. The W265G/M287V mutation of the P-protein W-hole impairs STAT1–STAT2 antagonism and also abolishes P-mediated inhibition of STAT3 nuclear accumulation, consistent with defective targeting of activated STAT1-containing heterodimers, including STAT1–STAT2 and STAT1–STAT3 complexes^40,43^. Duvenhage virus (DUVV), in which the residue corresponding to Trp265 of P protein is changed to Gly, is defective for STAT1–STAT3 heterodimer antagonism compared with RABV^40^.

### P-CTD interacts with the NTD of pY-STAT1 using the site C

P-CTD that binds to the interface of the core regions of AB protomers of pY-STAT1 also interacts with the NTD of pY-STAT1 protomer C (Fig. 2d). At the pY-STAT1-C-NTD/P-CTD interface, the region spanning from β-sheet β2 to helix α2, as well as the region between helices α5 and α6 of the P-CTD interact with a pY-STAT1-NTD. This interaction involves residues Arg218, Phe223, Tyr225, Gln229, Lys273, Leu276, and Leu277 of P-CTD, residues Leu21, Asn93, Phe94, Asp97, Gln100, Met103, Ile104, Ser107, and Glu111 of pY-STAT1-NTD (Fig. 2d, Extended Data Fig.4G, H). Notably, residues Leu276 and Leu277 were previously predicted by NMR analysis using full-length U-STAT1 to interact with either the NTD or C-terminal part^45^; the present data confirm interaction with the NTD in the context of the STAT tetramer. The positively charged side chain of Arg218 of the P-CTD is presumed to form salt bridges with the negatively charged side chain of Glu111 in pY-STAT1 (Fig. 2d, Extended Data Fig.4G). The side chains of Leu276 and Leu277 of the P-CTD interact with the hydrophobic region formed by Leu21, Met103 and Ile104 of pY-STAT1 (Fig. 2d, Extended Data Fig.4H). As P-CTD (X chain) binding site is on the opposite side of the dimerization interface of the NTD (C protomer) (Fig. 1e), the P-CTD does not interfere with this dimerization (AC dimer of NTDs) of pY-STAT1. The fact that one of the P-CTD protomer (X chain) interacts with pY-STAT1 protomers A, B, and C indicates that the symmetrically related P-CTD (protomer Y) interacts with the pY-STAT1 protomers C, D, and B: even though the EM-map around the P-CTD Y protomer is not clear. As a result, P-CTD recognizes both NTD-mediated tetramerization of pY-STAT1 and the core interactions induced by pY-SH2 dimerization, thus the 4:2 complex is extensive and highly stabilized to inhibit multiple functions of STAT1.

### The P-CTD-binding site in pY-STAT1 overlaps with the importin- and DNA-binding sites

The binding sites of P-CTD to the protomers A and B of pY-STAT1 are located in the STAT1 DBD in the core regions. It has been reported that importin α5 binds to pY-STAT1, but not to U-STAT1, where it is proposed to recognize dimerization in STAT1 upon phosphorylation rather than directly recognizing phospho-Tyr701, prior to transporting pY-STAT1 into the nucleus^47–49^. P-CTD interacts with the region around residues 407–413 of STAT1, which has been reported to contain the dimer-specific nuclear localization sequence (dsNLS, specifically residues Leu407, Lys410, and Lys413)^47–49^ (Fig. 2e). The interaction between pY-STAT1 and importin α5 also requires the NTD of STAT1 in addition to the dsNLS in the DBD suggesting the importance of considering a tetrameric form of pY-STAT1^49^.

The binding sites of P-CTD also overlap with the residues of the DNA binding site (Fig. 1i, j). This provides a clear explanation for the previous observations from cellular and biochemical assays that the binding between pY-STAT1 and DNA is inhibited in the presence of RABV or P protein/P-CTD^41,45^. Fluorescence polarization (FP) was used to determine the *K*_D_ for the interaction of pY-STAT1-core and GAS DNA (16 nM) (Fig. 3a, c) and compared to that of between pY-STAT1-core and P-CTD (194 nM, below). Among the residues of STAT1 DBD that interact with P-CTD, Gln340 (main chain amide), and Lys413 (side chain) from interface A (Fig. 2a,b), and side chains of Gln340 and Lys410 from the interface B (Fig. 2a, c, Extended Data Fig.4F), are used to recognize DNA ^7^.

**Fig. 3.**
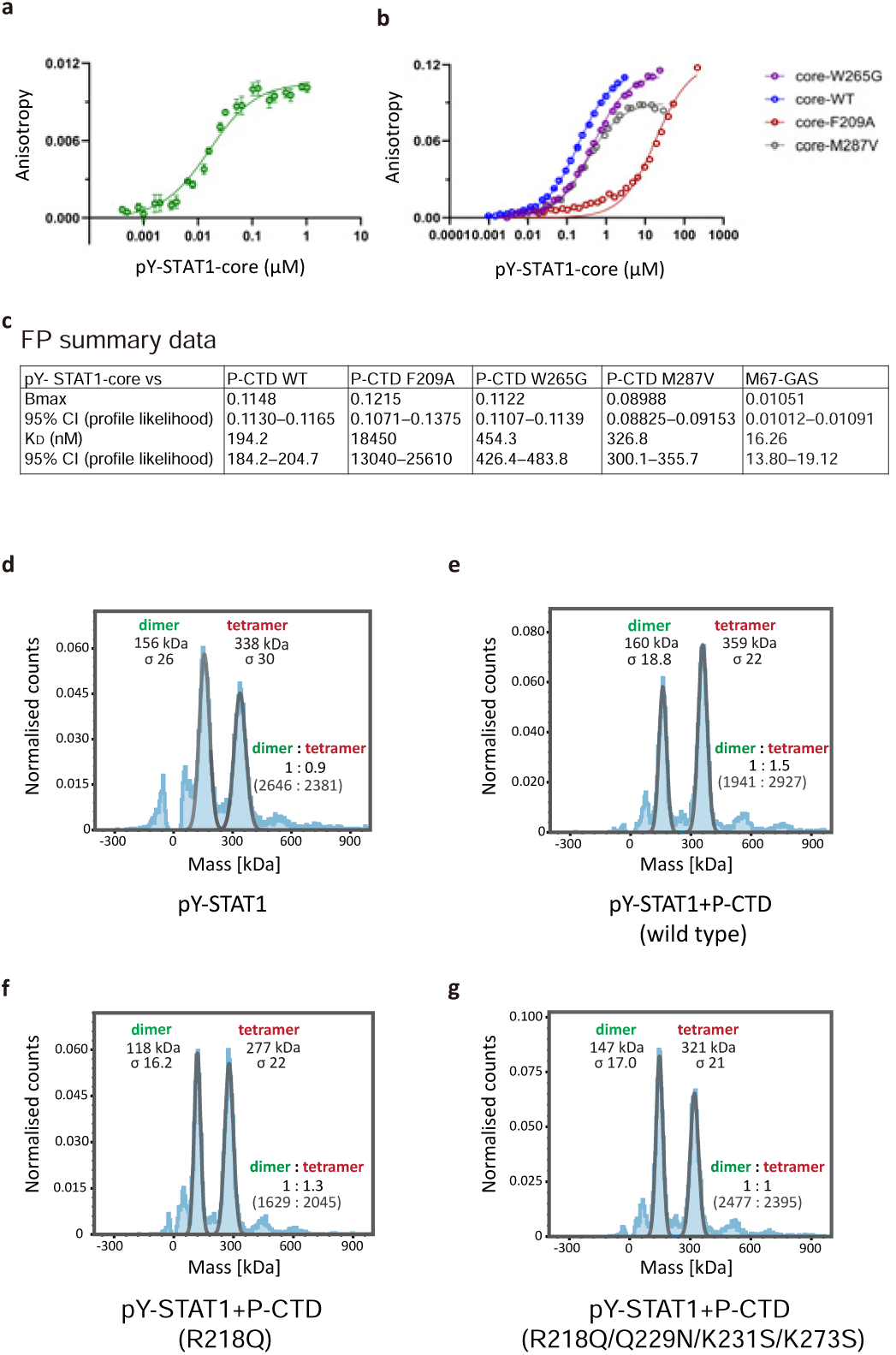
Characterization of the binding of DNA and of the P-CTD to the DBD of pY-STAT1. **(a)** Fluorescence polarization analysis of fluorescein labeled M67-GAS and pY-STAT1-Core. (**b)** Fluorescence polarization analysis of pY-STAT1 core and FlAsH-EDT_2_–labeled P-CTD (WT, blue; F209A, red; W265G, purple; M287V, gray). Data in **(a)** and **(b)** are shown in anisotropy value as mean ± SEM from three independent experiments (*n* = 3) for each combination. Data were analyzed with GraphPad Prism using the one site specific binding least squares fit function to calculate the *B*max and *K*_D_ values. **(c)** Data in **a, b** were analyzed with GraphPad Prism using the one site specific binding least squares fit function to calculate the *B*max and *K*_D_ values; mean *B*max and *K*_D_ values with SEM are summarized in the table. **(d) – (g)** Mass photometry analysis of the oligomeric state of pY-STAT1. Counts of dimers and tetramers detected by mass photometry are shown in brackets. **(d)** Oligomeric states of pY-STAT1 alone. **(e)** Oligomeric states of pY-STAT1 in the presence of wild-type P-CTD. **(f)** Oligomeric states of pY-STAT1 in the presence of P-CTD R218Q. **(g)** Oligomeric states of pY-STAT1 in the presence of P-CTD R218Q/Q229N/K231S/K271S.

### The binding of P-CTD affects the equilibrium of pY-STAT1 dimer–tetramer dissociation

pY-STAT1 is in equilibrium between tetramer and dimer (and monomer). The dissociation constant (*K*_D_) value of tetramer into two dimers in solution was reported to be 100 nM by analytical ultra centrifugation^50^. As the EM structure demonstrated that P-CTD recognizes tetramerization using the binding site C for pY-STAT1-NTD in addition to the pY-STAT1 dimer-recognition binding sites A and B, we monitored the effect of the P-CTD on the tetramer–dimer equilibrium of pY-STAT1 using mass photometry^51^. The ratio of dimer:tetramer of pY-STAT1 is 1:0.9 at a concentration of 53 nM (Fig. 3d). When P-CTD was introduced at 53 nM, the ratio of dimer:tetramer of pY-STAT1 increased to 1:1.5, Fig. 3e). The effect of P-CTD on stabilizing the pY-STAT1 tetramer was markedly reduced when residues of the P-CTD involved in site C were mutated, namely R218Q/Q229N/K231S/K273S, shifting the dimer:tetramer ratio to 1:1. The single P-CTD R218Q mutation had a modest impact (dimer:tetramer, 1:1.3, Fig. 3f, g). Analysis using the circular dichroism (CD) spectroscopy, which showed that no structural disruption was introduced by these mutations, supported that the mutations have specifically antagonized the pY-STAT1-NTD interaction (Extended Data Fig.5). The mass photometry experiment clearly demonstrated that P-CTD stabilizes the pY-STAT1 tetramer to inhibit dissociation into a dimer or inhibit the transition into a functional tetramer to cooperatively bind to DNA (Fig. 3)^16^. This effect of stabilizing the compact pY-STAT1 tetramer is expected to be enhanced for intact P protein as it dimerizes via the DD (Fig. 1b).

### Conservation of binding sites in STAT and P proteins

The amino acid sequence of STAT1 from mammals is highly conserved among mammals (Extended Data Fig.6). The residues that interact with P-CTD are completely conserved (Extended Data Fig.6). Consistent with this RABV has been shown to prevent the nuclear transport of STAT1 in mouse and bat cells^52,53^ suggesting that RABV P will be able to inhibit STAT1 signaling, and with the reported capability of RABV to infect many or all mammals, including the ability to cause disease in experimentally infected bats.

The amino acid sequence of human STAT1 exhibits 38% identity and 54% similarity with human STAT2, and 51% identity and 70% similarity with human STAT3 (Extended Data Fig.7), and its overall three-dimensional structure is well aligned with the reported structures of each of these STATs^6,7^ (Extended Data Fig.7). In region A of the STAT1 DBD, among the residues whose side chains are involved in P-CTD recognition, Leu383 is identical in human STAT2 and STAT3, whereas Gln340, His406, and Gln408 are not conserved. In region B, Asn381 and Leu383 are identical, while Gln340 and Lys410 are not. Residues involved in P-CTD recognition at region C of the STAT NTD showed lower conservation compared with those in regions A and B. Specifically, Leu21 and Glu111 are identical, whereas Asn93, Asp97, Gln100, Met103, and Ile104 are not conserved. From this sequence comparison, we cannot predict whether P-CTD will show preference for targeting among the human STAT1–STAT1, STAT1–STAT2, or STAT1–STAT3 dimers, or the key determinants for any such discrimination.

Among the CTDs of lyssavirus P proteins, the region encompassing residues ∼198–238 is the most conserved (Extended Data Fig.8). The eight amino acid residues of region A of which side chains are used for the STAT1 recognition are almost completely identical throughout the genus. Of the eight amino acid residues belonging to region B, half the residues are conserved including Pro245, Gly246, Arg249 and Met287, but the others that make contacts, Asn243, Trp265, Ala280, and Asn284 are not conserved. In the eight amino acid residues belonging to region C, most residues are conserved, although proteins of several lyssaviruses contain substitutions of Lys272 and Lys273 to the similar basic Arg residue.

### Interaction analyses of pY-STAT1 and P-CTD indicate site A preferentially binds to pY-STAT1

To investigate the contribution of side chains of P-CTD observed in the P-CTD-STAT1 structure to the interactions, fluorescence polarization (FP) assays were used to determine dissociation constants (*K*_D_) for binding between P-CTD variants (F209A [site A], W265G [site B], and M287V [site B]) and the pY-STAT1 core (Fig. 3b, c). Relative to the wild type (*K*_D_ = 0.19 µM)^16^, the F209A substitution resulted in a pronounced reduction in binding affinity (*K*_D_ = 18.5 µM), while the W265G and M287V substitutions caused only modest decreases (*K*_D_ = 0.45 and 0.33 µM respectively). To assess the structural integrity of the P-CTD mutants (F209A, W265G, and M287V), we used CD spectroscopy. The CD spectra of all variants were similar to that of wild-type P-CTD, indicating that the increased *K*_D_ values did not result from global structural distortion, but rather from the loss or weakening of direct side-chain-mediated interactions (Extended Data Fig.5).

Previously, transferred cross-saturation transverse relaxation-optimized spectroscopy (TCS-TROSY) NMR experiments were successfully performed for wild-type P-CTD with U-STAT1, but experiments could not be conducted with pY-STAT1 due to strong binding to the P-CTD^45^. We took advantage of the weak binding of the F209A mutant (Fig. 3b,c) to successfully perform TCS-TROSY measurements with N-terminal truncated pY-STAT1 (pY-STAT1-ΔN), which prevents dimerization/tetramerization via the N-terminal region (Fig. 4). P-CTD WT with U-STAT1-ΔN and the P-CTD F209A mutant with pY-STAT1-ΔN exhibited the same interaction effects at site A (Fig. 2b), accompanied by marked attenuation in the intensity ratios of the ^15^NH resonances of His203, Ala206, and Asp235. In contrast, interaction at site B (Fig. 2c) was observed only for the interaction of the P-CTD F209A mutant with pY-STAT1-ΔN). For the interaction between P-CTD WT and U-STAT1-ΔN, no significant attenuation was detected in the intensity ratios of the ^15^NH or ^15^N^ε^H resonances of Trp265^45^. Upon binding to pY-STAT1-ΔN, the ^15^N^ε^H resonances of Trp265 of P-CTD F209A showed substantial attenuation. These results suggest that Trp265 in the P-CTD, and therefore site B, specifically interacts with pY-STAT1 but not with U-STAT1, and that U-STAT1 preferentially interacts with site A of the P-CTD. These data are consistent with the idea that the lack of detection of binding by the W-hole in previous structural analyses using U-STAT1^45^, but clear effects of W-hole mutants on antagonism of pY-STAT1-dependent signaling relate to roles of the W-hole in binding to pY-STAT heterodimer (pY-STAT1–STAT2)^43^. Mechanistically these differences suggest that site A makes an initial contact, followed by an induced fit of region B to obtain a high affinity complex in a pY-STAT1 dimer.

**Fig. 4.**
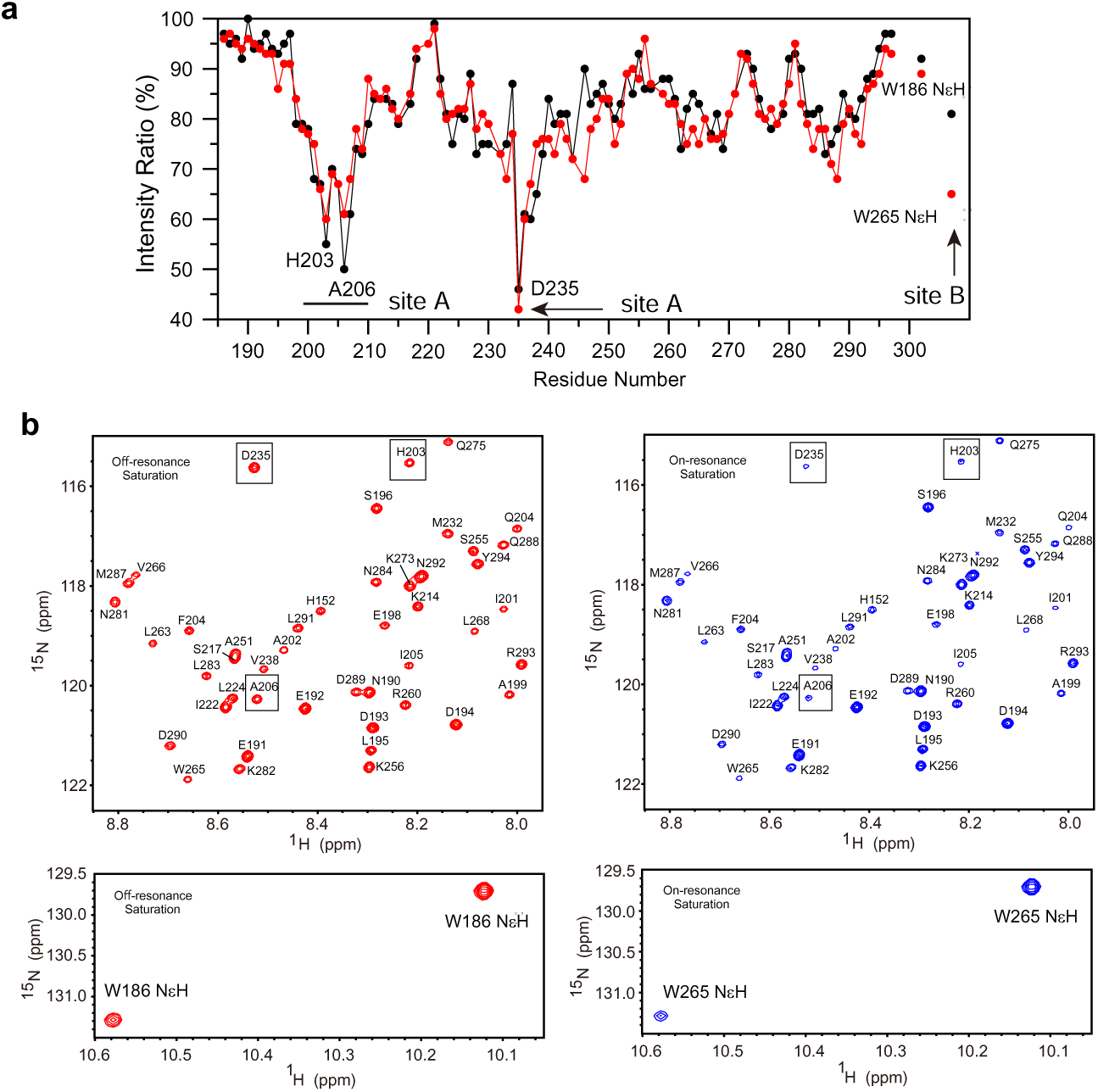
Trp265 of P-CTD shows an interaction with phosphorylated but not unphosphorylated STAT1. **(a)** Intensity ratios (%) of ^15^NH resonances from on- and off-resonance transferred cross-saturation ^15^N,^1^H transverse relaxation-optimized spectroscopy (TROSY) experiments of wild-type P-CTD with unphosphorylated N-terminally truncated STAT1 (U-STAT1-ΔN) (black circles) and F209A P-CTD with phosphorylated pY-STAT1-ΔN (red circles), at 298 K and pH 6.8. Under these solution conditions both U-STAT1-core and pY-STAT1-core show interactions with site A of the P-CTD constructs (Fig. 2b) with intense attenuation of the ^15^NH resonances of His203, Ala206 and Asp235. However, only pY-STAT1-core shows an interaction with site B (Fig. 2c) for F209A P-CTD where the Trp265 ^15^N^ε^H is attenuated. These results suggest a preferred interaction of site A of the P-CTD for unphosphorylated STAT1. **(b)** Spectral regions of the off- and on-resonance (left and right panels respectively) transferred cross-saturation TROSY of F209A P-CTD with pY-STAT1-core highlighting attenuation of the ^15^NH resonances Ala206 and Asp235 in the upper spectra and ^15^N^ε^H of Trp265 in the lower spectra.

### Region B mutations in P protein impair antagonism of the type I IFN (pY-STAT1–STAT2-dependent) but not the type II IFN (pY-STAT1–STAT1-dependent) pathway

To investigate which regions of the P-CTD may recognize and discriminate between the pY-STAT1 homodimer and the pY-STAT1–STAT2 heterodimer, we performed luciferase reporter gene assays to monitor the effects of mutations of sites A (F209A) or B (W255G or M287V) of the P-CTD on antagonism of the type I or type II interferon (IFN) signaling pathways by P protein (Fig. 5). IFN-γ–induced pY-STAT1-dependent signaling was monitored using a luciferase reporter gene plasmid with a GAS-containing promoter (pGAS-luc), while IFN-α–induced pY-STAT1–pY-STAT2-dependent signaling was monitored using a plasmid with an ISRE-containing promoter (pISRE-Luc). Consistent with previous reports, expression of wild-type P protein strongly inhibited both type I and type II IFN signaling compared with the control (GFP-fused RABV N protein)^36,45^. The inhibitory effect of P protein on both type I and type II IFN signaling was markedly reduced by F209A mutation. However, while Trp265 or Met287 mutation strongly impaired signaling by type-I IFN (STAT1–STAT2-dependent) as previously observed^43,45^, both mutants retained antagonistic function toward type II IFN signaling comparable to WT. These data suggest that sites A and B preferentially interact with STAT1 and STAT2, respectively, in the pY-STAT1–STAT2 heterodimer whereas binding of P-CTD to pY-STAT1–STAT1 is more dependent on site A.

**Fig. 5.**
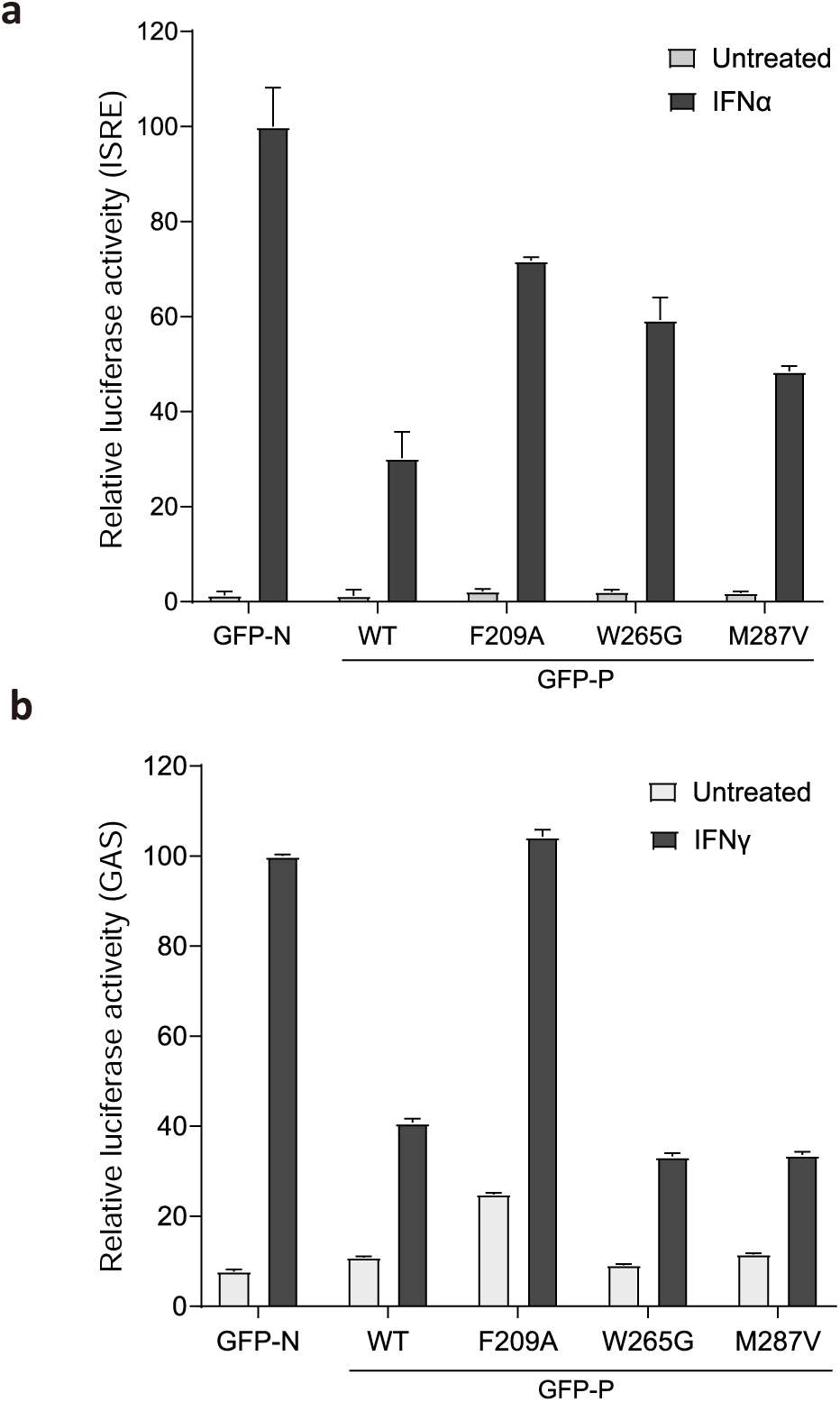
Analysis of the effects of mutations of site A or site B of the P-CTD on P protein-mediated antagonism of type-I and type-II IFN-dependent signaling. Cos-7 cells were co-transfected to express the indicated GFP-fused P-proteins or N-protein control, together with plasmids for luciferase reporter gene assays: (**a**) pISRE-luc (for type I IFN signaling) or (**b**) pGAS-luc (for type II IFN signaling), which express firefly luciferase under the control of an ISRE or GAS-containing promoter, respectively, and (**a**, **b**) pRL-TK plasmid that expresses *Renilla* luciferase constitutively (transfection control). Cells were treated 6 h post-transfection without or with (**a**) 1000 U of type I IFN or (**b**) 100μg/ml of type II IFN, and analysed by a dual luciferase assay 16 h post-treatment. Values for firefly luciferase were normalized to values for *Renilla* luciferase and calculated as fold change compared with values for GFP N-protein transfected cells treated with IFN; mean ± SEM (n = 3); data are from a single assay representative of 3 independent assays.

## Discussion

Many pathogenic viruses encode antagonist proteins that target and inhibit STAT proteins, thereby shaping viral pathogenesis and host tropism. However, the structural details of these interactions are generally poorly defined leaving molecular mechanisms of antagonism, which are important to understanding of immune evasion and its potential targeting for intervention, unresolved. Our structural analysis now reveals how the RABV P-CTD specifically recognizes the activated form of STAT1 through three distinct interfaces. Combined with biophysical and IFN-signaling analyses, our findings provide a mechanistic explanation for how the RABV P protein counteracts JAK–STAT-mediated antiviral immunity through multiple, coordinated mechanisms: inhibition of nuclear translocation, blockade of DNA binding, and a novel mechanism of restricting the conformational transition of the pY-STAT1 tetramer to prevent the formation of the extended DNA-binding state.

Our data, indicate that inhibition of pY-STAT1 nuclear import by P protein involves specific binding of the P-CTD to an importin α5-recognition site in STAT1 (including residues Lys410 and Lys413 in the DBDs of both A and B protomers), which is produced by dimerization of pY-STAT1. The pY-STAT1 dimer or tetramer is recognized by importin α5 with an interaction that is weaker than that of pY-STAT1 and P-CTD (pY-ΔN-STAT1 binds importin α5 with a *K*_D_ of 112 nM and P-CTD with a *K*_D_ of 37.6 nM; pY-STAT1 binds importin α5 with *K*_D_ of 191 nM and P-CTD with *K*_D_ of 18.1 nM)^16,49^ (Fig. 6). This suggests that the binding of P proteins to the pY-STAT1 dimer and/or tetramer will prevent importin interaction and active nuclear import of STAT1. In addition to this effect, the specific localization of different isoforms also contributes to inhibition of nuclear localization. Specifically, RABV P protein is expressed as at least five isoforms, the full-length P protein (known as P1) and P2–P5, which are N-terminally truncated forms of P1, produced by translation from internal in-frame start codons (Met20, 53, 69 and 83 for P2, P3, P4, and P5, respectively)^54^. Only P1 contains the L protein-binding site in the N-terminal region, and so is the only form that acts as the essential cytoplasmic polymerase cofactor^55^, but all isoforms contain the STAT1-binding P-CTD and DD, enabling roles in immune evasion. Notably the isoforms differ significantly in subcellular localization, whereby P1 and P2 are cytoplasmic due to the presence of a strong N-terminal nuclear export sequence (NES, residues 49 to 58), resulting in strong localization of P–pY-STAT complexes to the cytoplasm. The shorter isoforms P3–P5 lack the N-terminal NES and use other targeting sequences to variously localize between the cytoplasm, nucleus, nucleolus, and nuclear bodies, and to associate with cytoplasmic microtubules (MTs), where the latter can effect sequestration of STATs to MTs^36,38,41–43,53,56–64^. The binding to key residues of the DBD also appears to account for impaired binding of STAT1 to DNA by the P-CTD^45^, consistent with an intranuclear inhibition by the shorter isoforms^41^.

**Fig. 6.**
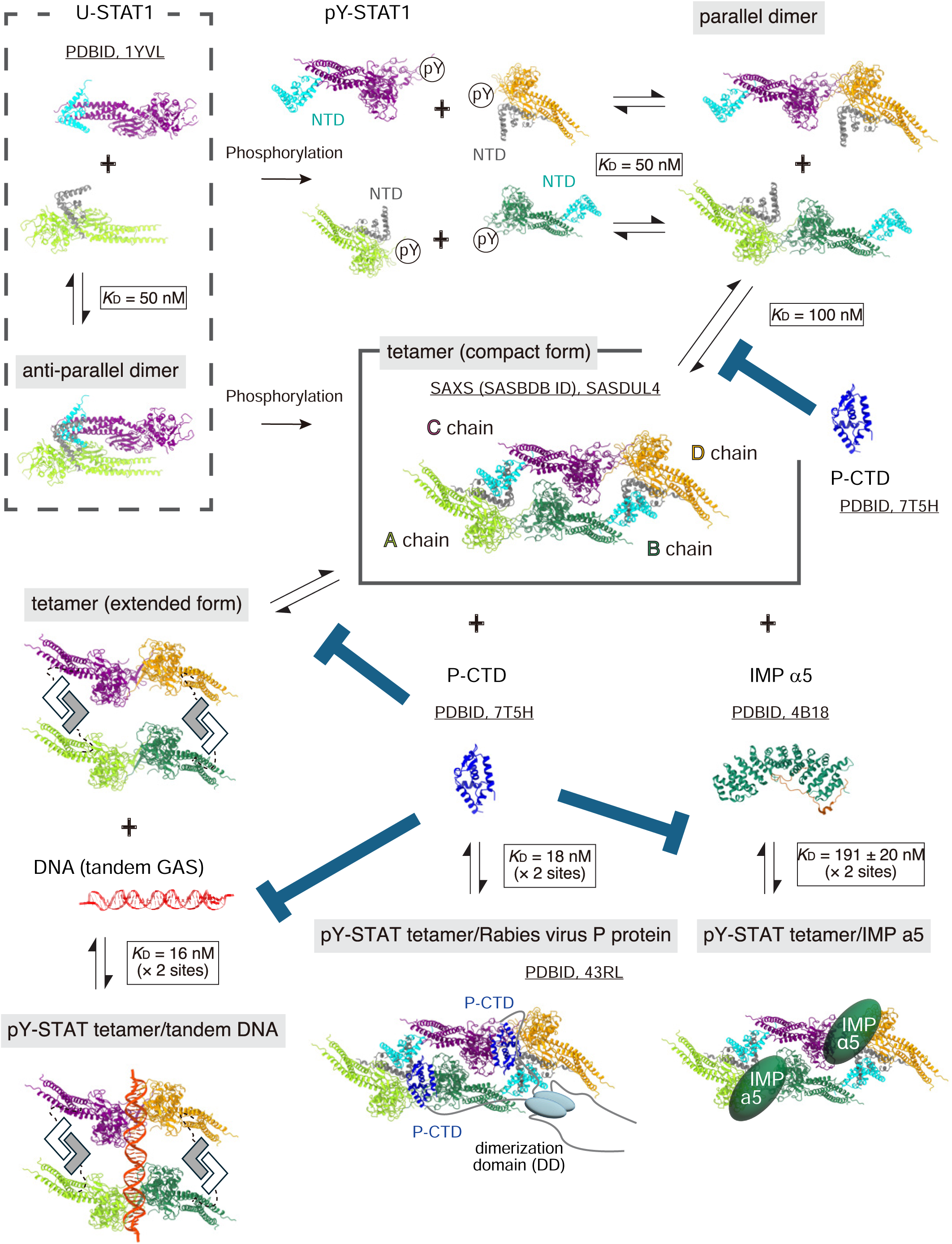
Equilibrium of STAT1 oligomerization and competitive interactions between host factors and P-CTD. STAT1 oligomerizes through tyrosine phosphorylation and NTD-mediated interactions (top). The STAT1-core region of four protomers (A–D chains) are colored in light green, green, purple, and orange respectively. The NTDs of A and D chains are colored in gray, and NTDs of B and C chains are colored in cyan. RABV-P-CTD (colored in blue) binds to the pY-STAT1 tetramer, forming a complex (bottom center). Because P protein forms a dimer via DD, two P-CTD molecules are expected to associate with a single pY-STAT1 tetramer. P-CTD inhibits the transition of the pY-STAT1 tetramer to its functional form (center left), as well as its binding to DNA (bottom left) and importin α5 (center right). The stoichiometry of the importin α5–pY-STAT1 interaction is 2:1^49^, suggesting that two molecules of importin α5 bind to a pY-STAT1 tetramer. Corresponding *K*_D_ values for each equilibrium and the PDB IDs of the referenced structures are indicated.

The equilibrium of STAT1 tetramerization in solution is complicated (Fig. 6). U-STAT1 likely forms an anti-parallel dimer with *K*_D_ of 50 nM^50^. This interaction is stronger than that of two isolated NTDs (*K*_D_ = 17–23 μM^50,65^), as additional regions such as the CCD of one molecule forms reciprocal interactions with the DBD of the other, as indicated by the crystal structure^66^. Upon the phosphorylation of Tyr701, two pY-STAT1 proteins interact with one another via the reciprocal pY–SH2 (*K*_D_ of 50 nM) to form a parallel dimer. Further, two pY-STAT1 dimers tetramerize with a *K*_D_ of 100 nM^50^. In the nucleus, pY-STAT1 dimer bind to single GAS sequences (*K*_D_ of 16 nM in solution, measured in the manuscript using pY-STAT1-core dimer, Fig.3a, c); this *K*_D_ shows stronger binding than that of P-CTD binding to the pY-STAT1-core dimer (*K*_D_ = 194 nM^16^). However, it is plausible that in full length P protein isoforms (all of P1–P5), the avidity for pY-STAT1 will be increased, enhancing the capacity to inhibit pY-STAT1-DNA binding, because all of the P isoforms harbor a DD and so will present dimers of the P-CTD. Thus, the inhibition of DNA binding, especially by nuclear populations of P3–P5 protein, is likely to contribute to antagonism^41^.

Using mass photometry, we also observed that P-CTD acts to stabilize the pY-STAT1 tetramer preventing dissociation into dimers; this also accounts for preventing transition into the NTD-tethered extended tetramer (Fig. 3d–g, 6), which was proposed as a functional form^16^. The finding that P-CTD inhibits conformational transition (and DNA binding) of the complex is also consistent with the observation that P protein inhibits the normal phosphorylation/dephosphorylation cycle of pY-STAT1 as a part of its antagonistic function, retaining pY-STAT1 in non-active complexes in IFN-treated cells expressing P protein or infected by RABV. Taken together, P-CTD plausibly hinders three functions of STAT1: nuclear import, DNA binding, and structural transition into the functional form (Fig. 6).

P-CTD has also been reported to interact with RABV N protein-RNA complexes^35,67^, promyelocytic leukaemia (PML) protein^68^, nucleolin^69^, and microtubules^58^. Notably, both N protein and pY-STAT1 can be co-immunoprecipitated together when co-expressed with P-CTD (but not without P-CTD), indicating that P-CTD can simultaneously bind and, thus, link these proteins^35^. These data indicate that not only free P protein but also N-RNA-bound P protein (which is involved in genome replication by linking P-associated L protein to the N-RNA) can bind to and inhibit pY-STAT1^35^. P-CTD binds to the C-terminal arm (CTARM) of N protein in the N-RNA complex and recruits L protein that is bound to the N-terminal region (NTR) of P1 (primary residues 1–19; additionally residues 51–87)^55,70,71^. The structural analysis in the current study supports the model that both free-unbound and N-RNA-bound P protein can bind to pY-STAT1 tetramers, because the reported binding site on the P-CTD for N-RNA (including residues Lys211, Lys214, Leu224, Arg260)^35^, does not overlap with any of the three pY-STAT1 binding sites identified (Fig. 7).

**Fig. 7.**
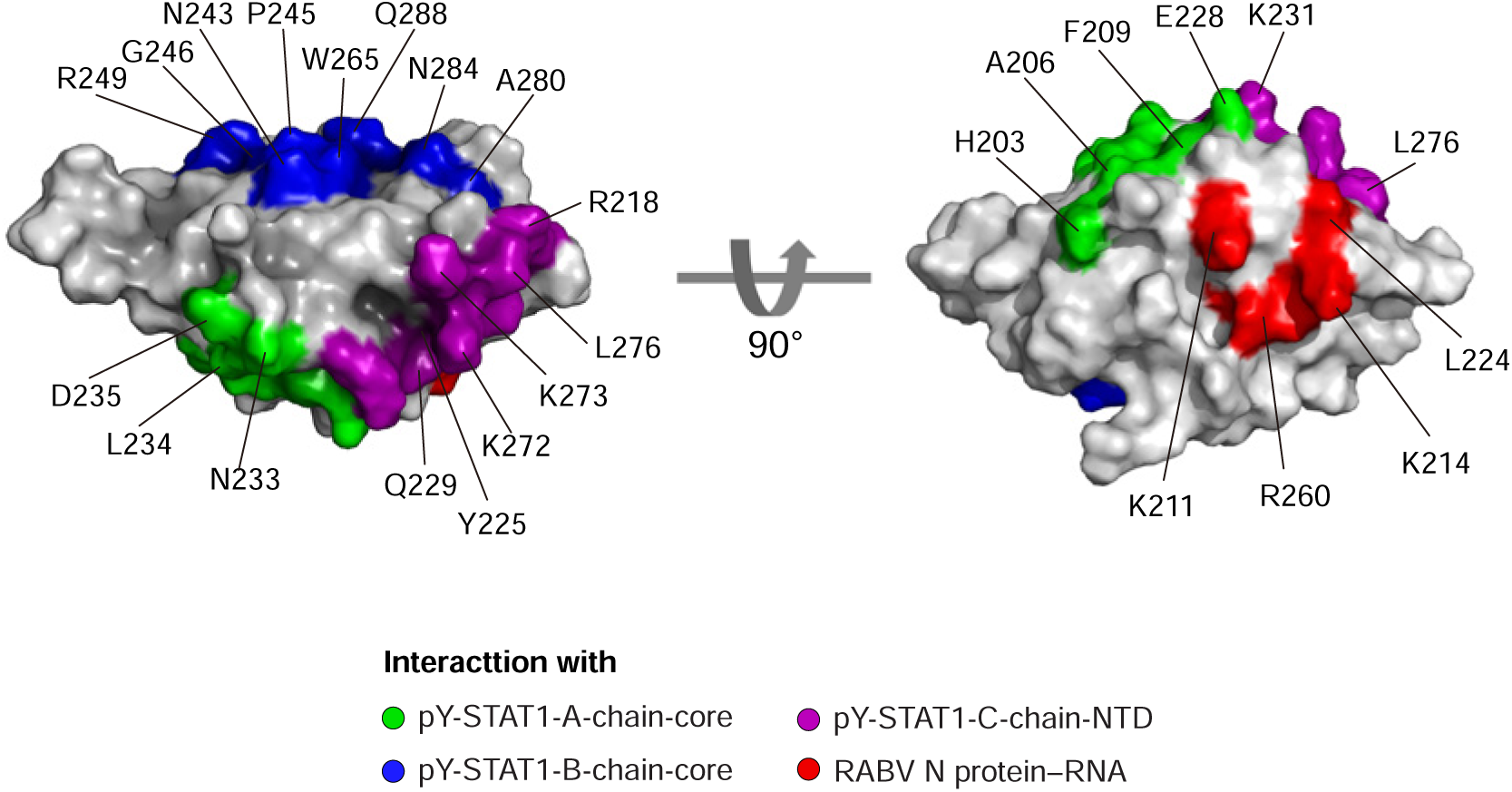
P-CTD surface representing pY-STAT1 and N binding sites. The binding site on the pY-STAT1 A chain core is shown in light green, that on the B chain core in blue, and that on the C chain NTD in purple. The site shown in red corresponds to the RABV N protein–RNA complex binding region proposed in previous studies^35^, which is located at a position distinct from the STAT1 interaction sites.

A substitution at Asn226 to His in the P-CTD was identified in Ni-CE, an attenuated RABV strain, compared to the parental strain Nish^62^. The side chain of Asn226 is located on the P-CTD surface proximal to Ser210, Glu228, and Gln229 (Extended Data Fig.4I). Among these, Glu228 is in site A and Gln229 is in site C, thus directly interacting with pY-STAT1. Notably, Ser210, along with Ser271, has been reported to be phosphorylated by host protein kinase C (PKC)^72^. Modifying the local environment around this region of P3 by introducing the Ni-CE mutation N226H, or a phosphomimetic mutation (S210D or S210E) result in defects in both nuclear accumulation and MT association, correlating with effects on phase separation activity of “pathogenic” P3^57,58,73^. Similarly, P proteins harboring the N226H substitution or phosphorylation of Ser210 may alter the local environment, modulating the interaction with pY-STAT1 although no direct interaction of Ser210 with pY-STAT1 was observed from our NMR experiments using the wild type P-CTD (the current and previous studies^45^).

Several structural studies on STAT proteins in complex with viral proteins have been conducted: the Sendai virus C protein with the NTD of human STAT1^28^; the poxvirus protein 018 with the core domain of STAT1^29^; the Zika virus and dengue virus nonstructural protein 5 (NS5) with the NTD and CCD of human STAT2^30,32^; and the rat cytomegalovirus E27 protein with the CCD of rat STAT2^74^. In these studies, truncated and unphosphorylated form of STAT molecules were used, although two protomers of the Sendai virus C protein recognize two NTDs of STAT1 (2:2 complex). In contrast, RABV P protein adopts a novel inhibitory strategy by targeting only the activated pY-STAT1 tetramer, which we show to be mediated through three distinct interaction surfaces. The use of these selective mechanisms, rather than targeting all cellular STAT1 including U-STAT1, presumably enables the efficient inhibition of activated pY-STAT1 using a limited number of P proteins. It also is expected to result in less disruptive effects on the physiology of infected cells, by specifically inhibiting cytokine-activated STAT1, an important aspect in the life-cycle of RABV, which requires the maintenance of neuronal networks for ingress and egress of the central nervous system^75^.

Other than revealing key aspects of virus-host immune evasion, the analysis of virus-host interfaces can reveal fundamental aspects of cellular biology, such as through the stabilization of cellular complexes for structural analysis. In our study^16^, we reported the tetrameric pY-STAT1 structure in complex with two DNA molecules. However, the NTD was not observed because of flexibility of the linker between the NTD and the CCD (residues 127–132). In this study, we were able to locate the NTD dimer because the position of the NTD is fixed through interaction with the P-CTD: a good example of utilizing a viral molecule as a probe to stabilize a protein complex (e.g., the tetramer of pY-STAT1).

Another prominent feature of P-CTD-mediated targeting of pY-STAT1 is that a twofold-related pY-STAT1-core dimer is recognized by a monomeric P-CTD, which itself lacks internal twofold rotational symmetry. In the pY-STAT1-core homodimer, corresponding regions of the two STAT1 DBDs are engaged independently by distinct surfaces of P-CTD, namely sites A and B (Fig. 2a). Notably, P-CTD selectively inhibits not only the activated pY-STAT1 homodimer but also activated pY-STAT1–STAT2 and pY-STAT1–STAT3 heterodimers, whereas it does not inhibit the pY-STAT3 homodimer^38,40^. These observations suggest that, in STAT1-containing heterodimers, with STAT2 or STAT3 may be engaged by either site A or site B, while the other site recognizes the DBD of STAT1. In fact, previous NMR analyses detected primarily site A when the interaction between monomeric U-STAT1-ΔN and P-CTD was monitored^45^. In the present study, both sites A and B were detected when P-CTD F209A was titrated with dimeric pY-ΔN-STAT1. Mechanistically these differences suggest that site A makes an initial contact with pY-STAT1, followed by an induced engagement of site B to obtain a high-affinity complex with the pY-STAT1 homodimer. In agreement with this interpretation, FP analysis using point substitutions at either site A or site B on the P-CTD surface showed that the reduction in affinity for pY-STAT1 core was more pronounced following mutation of site A (F209A) than site B (W265G or M287V) (Fig. 3b).

Luciferase reporter assays further indicated that substitutions at site B, W265G or M287V, selectively impaired antagonism of the type I IFN pathway, mediated by the pY-STAT1–STAT2 heterodimer, while having little effect on antagonism of the type II IFN pathway, mediated by pY-STAT1 homodimers and/or tetramers (Fig. 5). These findings suggest that, in the context of type I IFN signaling, site B contributes to recognition of STAT2, whereas site A primarily recognizes STAT1. Together, these observations suggest that P-CTD is tuned to discriminate among STAT homo- and heterodimers, prioritizing specific STAT targets through distinct interaction surfaces and affinities for a purpose.

Future molecular dissection using the approaches described here for multimeric STAT assemblies will further clarify these mechanisms, where comprehensive atomic-level information on how P-CTD recognizes the pY-STAT1 homodimer, pY-STAT1–STAT2 and pY-STAT1–STAT3 heterodimers may facilitate the rational design of antiviral reagents or attenuated vaccines for RABV. Extending this strategy to other viral antagonists of the JAK–STAT signaling pathway may lead to novel antiviral therapeutics or attenuated vaccines.

## MATERIALS AND METHODS

### Expression vector constructs and purification of human pY-STAT1

Tyrosine-phosphorylated STAT1 (pY-STAT1) was prepared by transforming BL21(DE3) TKB1 cells (Agilent Technologies) with pET21d plasmids encoding STAT1 with an N-terminal 10×His tag. Tyrosine phosphorylation at Tyr701 was achieved during expression using the BL21(DE3) TKB1 host system. pY-STAT1 was subsequently purified as previously described protocols^16,76^.

### Expression vector constructs and Purification of RABV P-CTD

For cryo EM analysis, the Nish-P-CTD (residues 186 – 297), protein was expressed using a plasmid containing Nish-P-CTD cloned into a modified pET-28(a) expression vector with an N-terminal 10×His tag, as previously described^44^. For mass photometry analysis, P-CTD was prepared using a plasmid encoding P-CTD with an N-terminal 10×His tag and a C-terminal tetracysteine (TC) tag, as previously described^16^. Based on TC-tagged P-CTD plasmid, two mutant constructs were generated: R218Q and R218Q/Q229N/K231S/K273S. P-CTD R218Q/Q229N/K231S/K273S mutant was generated by first introducing the R218Q mutation, followed by the Q229N and K231S mutations to produce the R218Q/Q229N/K231S mutant, and subsequently introducing the K273S mutation. The primer sequences for each mutants are as follows: P-CTD R218Q: 5′-CCTTCTCAGTCTTCAGGGATATTCTTG-3′ (forward) and 5′-CTGAAGACTGAGAAGGAAACTTGTACTTC-3′ (reverse); P-CTD Q229N/K231S: 5′-TGAGCATGAACCTTGATGATATAGTTAAGG-3′ (forward) and 5′-AGTTCTCAAAATTATACAAGAATATCCCTG-3’(reverse); P-CTD K273S: 5′-ATTCTAAGAGCTTCCAATTGTTAGTCGAAGCCAACAAG-3′ (forward) and 5′-ATTGGAAGCTCTTAGAATTGGCCAGAGCGACC-3‘(reverse). All constructs were subsequently purified according to previously reported procedures^44^.

### Cryo-EM sample preparation and data acquisition

pY-STAT1/P-CTD complex was prepared by mixing pY-STAT1 and P-CTD with a molar ratio of 1:1.3 (pY-STAT1: P-CTD). The sample was purified by size-exclusion chromatography (SEC) using a Superose 6 increase 10/300 column (Cytiva, USA) with EM buffer (10 mM HEPES-NaOH, pH 7.4, 150 mM NaCl, 5% glycerol, 2 mM MgCl₂, 0.1 mM EDTA, and 2 mM DTT). The peak fraction, at a concentration of 0.2 mg/mL, was used for grid preparation. 2.5 microliters of the sample were applied onto glow-discharged (3 min, 10 mA on both sides using PIB-10 (Vacuum Device Inc.)) 300-mesh R1.2/1.3 Quantifoil Au grids. Grids were blotted for 15 s with a blotting force of 5 at 100% humidity and 4°C, then plunge-frozen in liquid ethane using a Vitrobot Mark IV (Thermo Fisher Scientific, USA).

The data were collected on a Krios G4 cryo-transmission electron microscope (Thermo Fisher Scientific, USA) equipped with a thermal field emission electron gun operated at 300 kV, GIF-Biocontinuum energy filter (Gatan, USA) with a 20 eV slit width., and a K3 direct electron detector camera (Gatan, USA) at the Center for Research and Education on Drug Discovery, Faculty of Pharmaceutical Sciences, Hokkaido University. Data collection was carried out using EPU software (Thermo Fisher Scientific, USA) at a nominal magnification of ×130,000, corresponding to a pixel size of 0.67 Å, with a defocus range of –0.8 to –1.8 (dataset 1), –0.8 to –1.4 (dataset 2), and –0.8 to – 1.8 μm (dataset 3). Micrographs were acquired as movies with a total exposure time of 1.5 s, a total dose of 52.0 (dataset 1), 56.9 (dataset 2), and 52.1 (dataset 3) electrons per Å^2^, fractionated into 50 frames. A total of 3,605 (dataset 1), 19,162 (dataset 2), and 4,571 (dataset 3) movies were collected and employed for the subsequent steps of image processing.

### Image processing, 3D reconstruction, and structure refinement

The cryo-EM data for the pY-STAT1−P-CTD complex were processed using cryoSPARC v4.4.1^77^. Following the motion correction and gain normalization with the “Patch motion correction” algorithm, the contrast transfer function (CTF) was estimated with the “Patch CTF Estimation” job. Images with poor CTF fits were excluded, resulting in 3,605 (dataset 1), 19,162 (dataset 2), and 4,571 (dataset3) images. Initial particle selection was performed using a simple shape-based method using a “Blob Picker” job, and the particles, resulting in 581,276 particles (dataset 1), 2,943,912 particles (dataset 2), and 722,719 particles (dataset3) being extracted after downscaling to a pixel size of 2.73 Å. After particle extraction, 490,098 particles (dataset 1), 2,547,574 particles (dataset 2), and 669,590 particles (dataset3) were selected after three rounds of 2D classification. Initial models were then generated by an “Ab-initio Reconstruction” job followed by one round of heterogenous refinement applying *C*1 symmetry. After another round of 3D classification using heterogeneous refinement, three classes containing pY-STAT1 dimer with well-defined density for pY-STAT1 N-terminal domain (pY-STAT1-NTD) and two classes containing pY-STAT1 tetramer were selected. Each particle set was subjected to an “Ab-initio Reconstruction” job to remove junk particles. For the dimers, the particles were re-extracted with a pixel size of 0.99 Å. After alignment using the “Non-uniform Refinement” job, a “3D classification” job was performed, focusing on pY-STAT1-NTD and P-CTD without alignment was performed to obtain the three classes that clearly showed the pY-STAT1-NTD and P-CTD. The final map of the dimer, comprising 328,639 particles, was reconstructed using “Non-uniform Refinement” jobs, achieving a resolution of 3.10 Å (Calculation of masked FSC curves were carried out using RELION^78^, Extended Data Fig.2). For the tetramers, “3D classification” focusing on P-CTD was performed, leading to a class with two P-CTD bound to the pYSTAT1 tetramer. Particles in this class were selected and re-extracted with a pixel size of 0.99 Å, then a final map of the tetramer comprising 56,425 particles was reconstructed with “Non-uniform refinement” at a resolution of 3.47 Å (RELION^78^, Extended Data Fig.2).

The local resolution of the final maps was calculated using the “Local Resolution Estimation” job. The image processing workflow is depicted in Extended Data Fig.1. Figures related to data processing and reconstructed maps were prepared with UCSF Chimera X (version 1.9)^79^.

### Cryo-EM model building and refinement

For atomic model building, the cryo-EM maps were trimmed to remove unnecessary regions outside the molecules using the software Servalcat^80^. For model building of the pY-STAT1–P-CTD complex, the reported structures of the STAT1 (PDB 43QG)^7,16^ and P-CTD (PDB 43RL)^57^ were used to fit into the cryo-EM map in UCSF Chimera X (version 1.4) ^79^. The initial structural model was then subject to iterative model building using Coot (v0.9.6)^81^ and real-space refinement using Phenix (v1.8.2)^82^. A model-map Fourier shell correlation was calculated using the criterion of 0.5. Protein Data collection, data processing, and model-building statistics are summarized in Supplemental Table S1.

### Nuclear Magnetic Resonance spectroscopy

NMR data were collected at 25 °C on a Bruker 700 MHz Avance IIIHD spectrometer equipped with a triple-resonance cryoprobe. For transferred cross-saturation transverse relaxation-optimized spectroscopy (TCS-TROSY) experiments^83–85^ 500 µM uniformly labelled ^2^H,^15^N-labeled wild-type P-CTD or F209A P-CTD were mixed with GB1-tagged N-terminal domain truncated (ΔN) unphosphorylated or phosphorylated STAT1 respectively, dissolved in 50 mM Na_2_PO_4_, 100 mM NaCl and 1 mM DTT, pH 6.8 in 90% ^2^H_2_O/10% H_2_O. Prior to mixing, the P-CTD was kept in the same buffer for 8 hrs at room temperature and then for at least 24 hrs at 4 °C to allow amide exchange to reach equilibrium. Under these solution conditions wild-type P-CTD binds weakly to unphosphorylated STAT1, but too strongly to phosphorylated STAT1 for successful TCS-TROSY^45^. However, the mutant F209A P-CTD shows weak binding to phosphorylated STAT1 enabling TCS-TROSY. The TCS-TROSY experiments used a 2D ^15^N,^1^H TROSY-HSQC pulse scheme and WURST ^1^H-saturation pulse (15 ms, 2800 Hz bandwidth)^86^. Data were acquired with 25% NUS with interleaved rows for on- and off-resonance saturation, spectral widths of 12 ppm in ^1^H (2048 data points) and 27 ppm in ^15^N (512 data points). The WURST saturation of the aliphatic protons of P-CTD was 2 s with a saturation frequency set at 0.9 ppm for on-resonance and -50 ppm for off-resonance. Each TCS-TROSY experiment was acquired in 16 h with 72 scans per row and a recycle time between scans of 1 s. Spectra were reconstructed with SMILE^87^, processed using NMRPipe^88^ and analysed in NMRFAM-SPARKY^89^.

### Fluorescence polarization assays

To assess the binding affinity of pY-STAT1-core for M67-GAS, a DNA probe (5’ ACAGTTTCCCGTAAATGCAA-3’) labeled with 6-FAM™ (Sigma-Aldrich, Germany). 6-FAM - labeled single-stranded DNA was mixed with the non-labeled complement sequence (5’ TGCATTTCAGGGAAACTT 3’) at a 1:1 molar ratio in TE buffer [10mM Tris-HCl (pH7.5), 1mM EDTA]. The solution was heated to 90 ℃ and then allowed to cool slowly to room temperature for DNA annealing. Two independent concentrations of pY-STAT1-core were prepared, and each was diluted twofold on a black flat-bottom polystyrene NB microplate (Corning, USA). In total, 24 wells of different concentrations of pY-STAT1-core were prepared, and 6-FAM-modified M67-GAS was added to each well to a final concentration of 14 nM. The final volume per well was adjusted to 100 μl with buffer [10 mM HEPES (pH 7.4), 150 mM NaCl, 1 mM DTT, and 10% glycerol]. A control lacking pY-STAT1-core was also prepared and used for gain adjustment. Fluorescence polarization measurements were immediately recorded with a SpectraMax i3x plate reader (Molecular Devices, USA), and the assay was repeated in triplicate. One-site specific binding least-squares fit curves were plotted with software Prism 10.4.2 (GraphPad, USA).

### Assessment of oligomer formation by mass photometry

Measurements were performed on a TwoMP mass photometer (Refeyn Ltd, UK). Freshly purified pY-STAT1 was analyzed both alone and in combination with each P-CTD variant. For complex formation, pY-STAT1 and P-CTD were mixed at a 1:1 molar ratio in the MP buffer (10 mM HEPES-NaOH, pH 7.4, 150 mM NaCl, 5% glycerol, and 1 mM DTT). All samples were diluted to 106 nM in the MP buffer just before the measurement. Prior to the measurements, a coverslip from the MassGlass™ UC – Sample Prep Kit (Refeyn Ltd) was mounted onto the mass photometer, and a gasket from the same kit was placed on top. A gasket well was filled with 10 µl of the MP buffer, 10 µl of the sample were added, and the adsorption of biomolecules was monitored for 120 seconds using the AcquireMP software (Refeyn Ltd, Version 2025 R1). For converting the measured ratiometric contrast into molecular mass, BSA (66kDa, MARUWA F&B, Japan), human immunoglobulin G (150kDa, MARUWA F&B, Japan), thyroglobulin (660kDa, MARUWA F&B, Japan) were used for calibration. All mass photometry movies were analyzed using DiscoverMP (Refeyn Ltd, Version 2025 R1). All samples were measured three times.

### Circular Dichroism spectrophotometry

J725 CD spectrometer (Jasco) was used for spectrophotometry. P-CTD WT and variants were adjusted to 15μM in 10 mM HEPES-NaOH, 150 mM NaCl, pH 7.4 at 25°C using a quartz cuvette (0.1 cm path length). For each protein, eight scans were recorded, averaged, and subtracted from the eight averaged buffer scans. Mean residue ellipticity (MRE) (deg.cm2.dmol-1) was calculated using MRE = *θ*.MRW/10.*l*.*c*, where *θ* is the ellipticity (millidegrees), *l* is the pathlength (cm), *c* is the protein concentration (mg/mL), and MRW is the Mean Residue Weight calculated as MRW = *M*r/(*N*–1), where *M*r is the *M*_W_ of the protein (Da) and *N* is the number of residues.

### Luciferase reporter gene assays

For the type-I and type-II IFN signaling assays, confluent monolayers of Cos-7 cells, grown in high-glucose DMEM, supplemented with 10% foetal calf serum, were co-transfected (Lipofectamine 2000, Thermo Fischer Scientific) with pEGFP-C1 constructs encoding a GFP-fused N-protein of the Challenge Virus Standard (CVS) RABV strain (non-STAT-1 targeting control as previously^42,43,64^), or GFP-fused Nish P proteins corresponding to those used in the structural analysis, together with the pRL-TK plasmid (Promega; encoding constitutively expressed Renilla luciferase) and either pISRE-Luc (Stratagene, for type-I IFN signaling assays) or pGASluc (a kind gift of Danielle Blondel, Institut de Biologie Intégrative de la Cellule, Gif-sur-Yvette, France, for type-II IFN signaling assays). pISRE-Luc and pGASluc, express firefly luciferase under the control of an ISRE-containing promoter or GAS-containing promoter, respectively. At 6 h post-transfection, cells were treated with or without 1000 U/mL recombinant human IFNα (PBL Interferon Source) or 100ug/ml recombinant human IFNγ (Peprotech) for 16 h, as previously described^90^. Cells were lysed in passive lysis buffer (Promega) and luciferase activity measured by dual luciferase assay as previously^42,43,90^ in 96-well plates using a CLARIOstar plate-reader (BMG).

## Author contributions

Conceptualization: A.S., G.W.M., P.R.G., and T.O. Methodology: M.M., A.S., T.N., C.T.D., Y.T., F.Y., S. C., Y.I., K.W., S.I.-I., N.Y.M.H., T.A., M.S., S.K.,Y.S., G.W.M., P.R.G., and T.O. Investigation: M.M., A.S., T.N., C.T.D., Y.T., F.Y., S. C., Y.S., and T.O. Visualization: A.S., M.M., Y.S., and T.O. Supervision: G.W.M., P.R.G., Y.S., and T.O. Writing—original draft: A.S., Y.S., and T.O. Writing—review and editing: G.W.M., P.R.G., and T.O.

## Data and materials availability

The cryo-EM density maps and corresponding atomic models have been deposited in the Electron Microscopy Data Bank (EMDB) and the Protein Data Bank (PDB) with accession numbers EMD-82067 and 43QG for the dimer and EMD-82102 and 43RL for the tetramer. All data needed to evaluate the conclusions in the paper are present in the paper and/or the Supplementary Materials.

