## Supplemental Figure for "Rabies virus antagonizes interferon signaling by targeting phosphorylated STAT1 tetramers"

Supplementary Figure S1

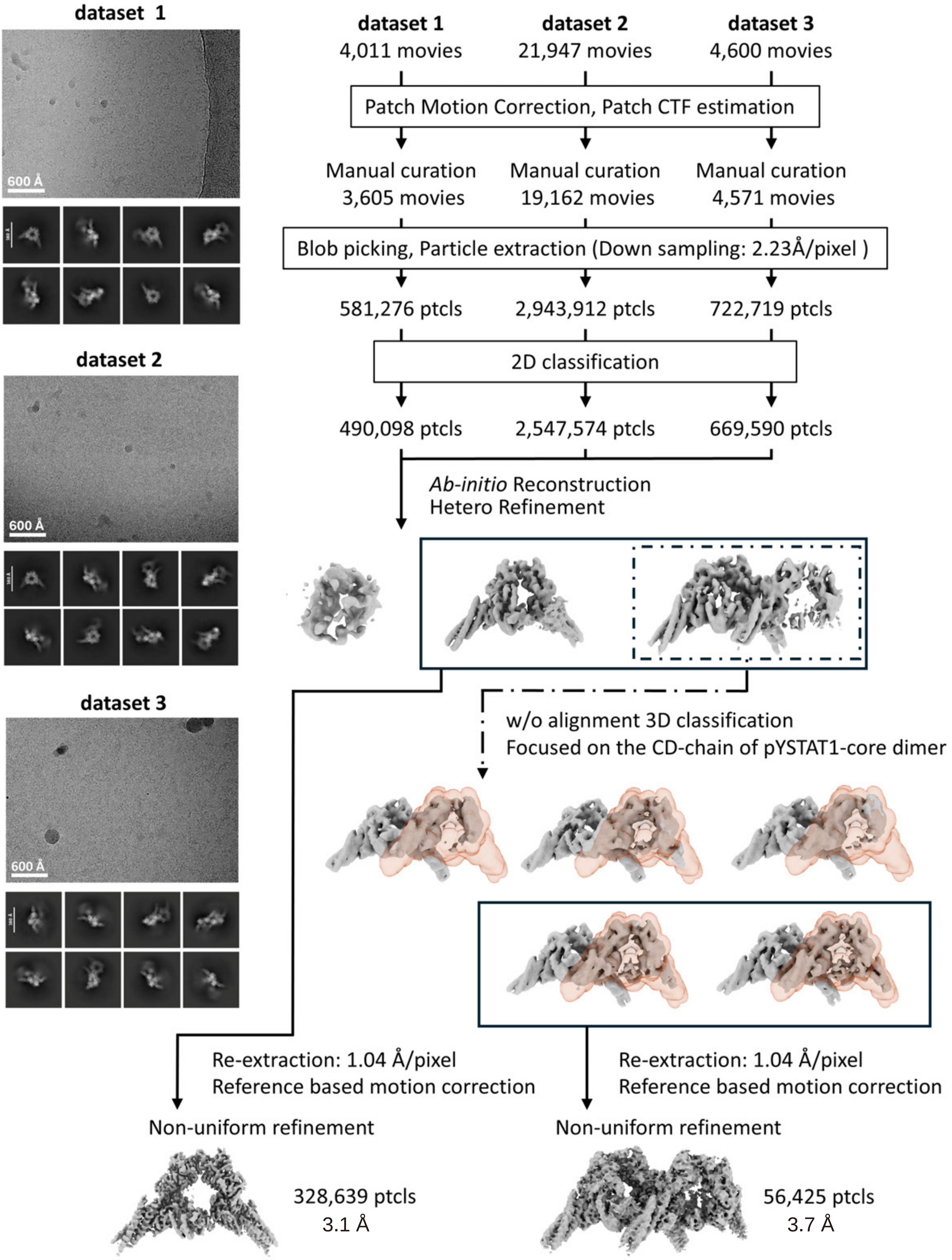

Supplementary Figure S2

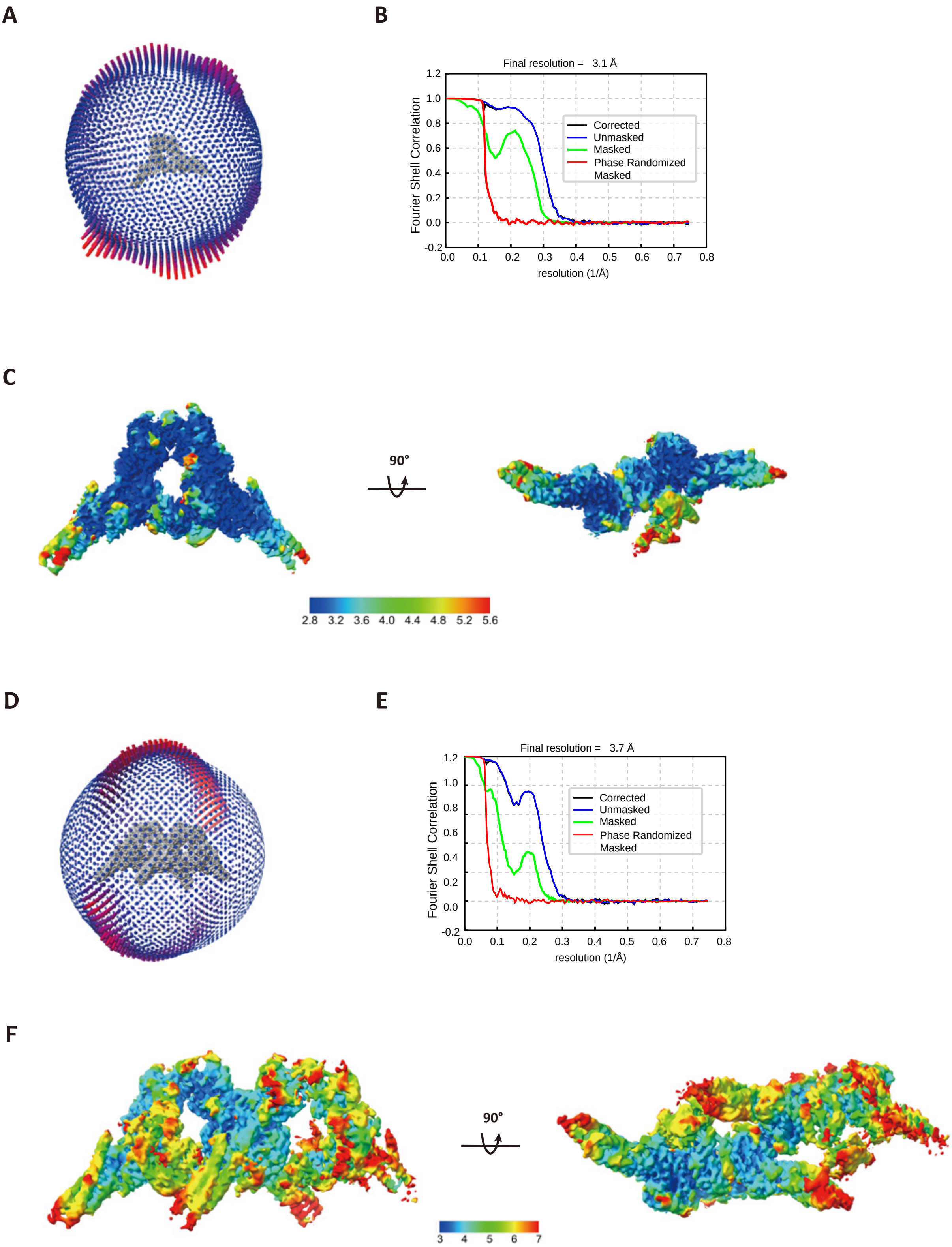

Supplementary Figure S3

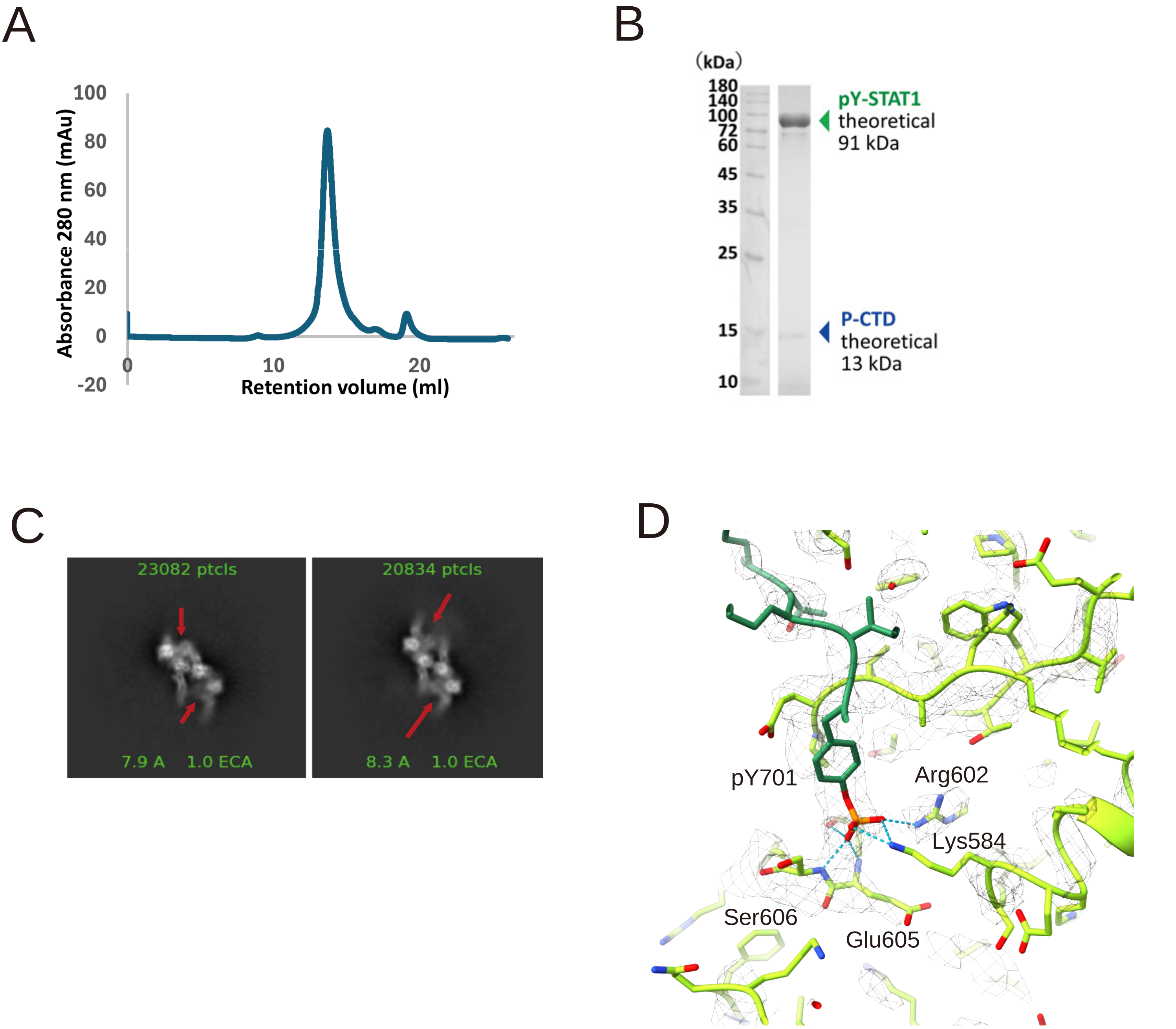

### Supplementary Figure S4

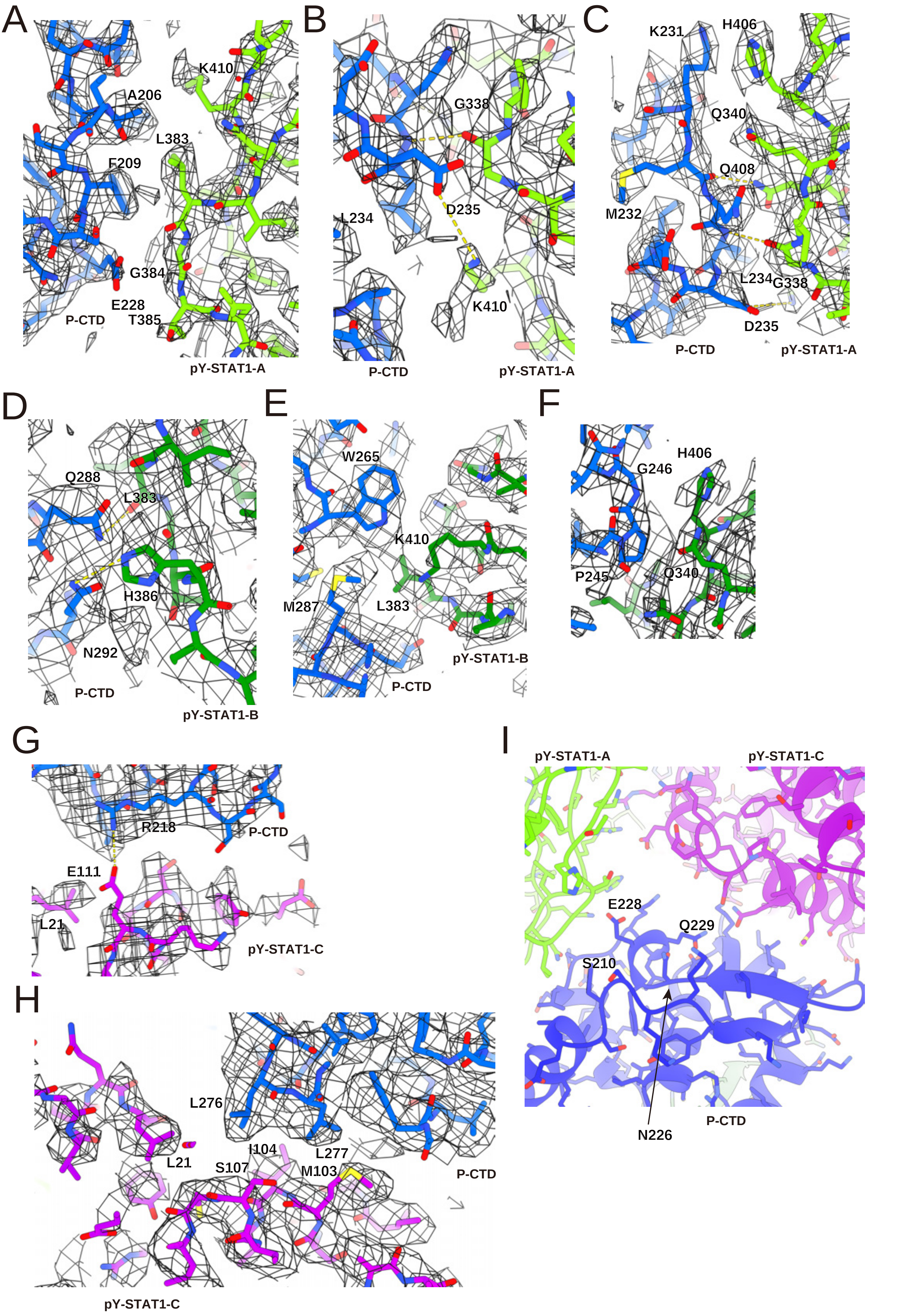

Supplementary Figure S5

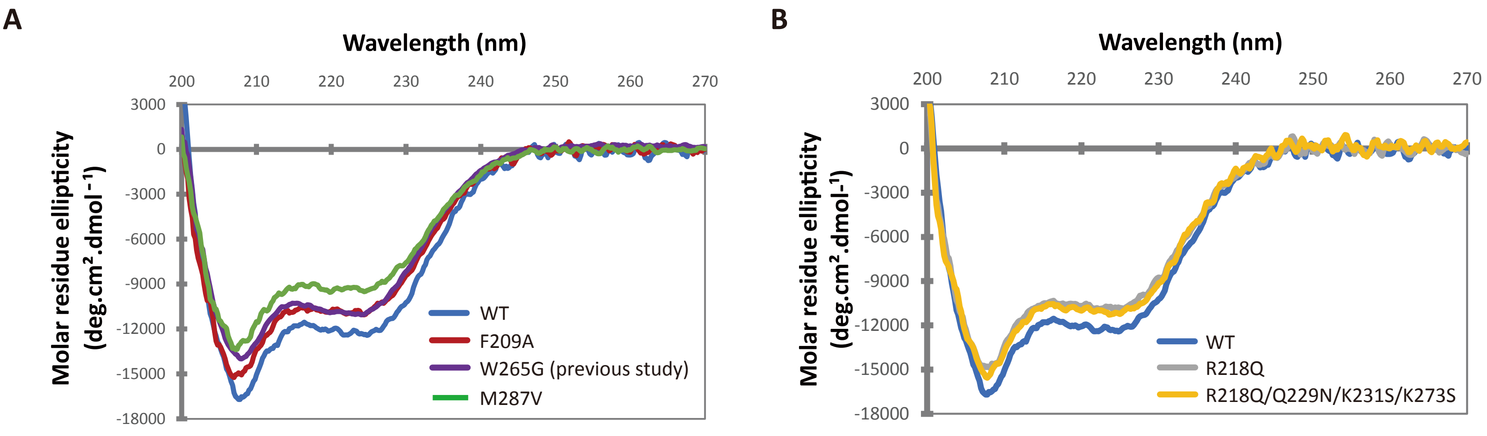

### Supplementary Figure S6

```
Human             ............................................................
Bat               ............................................................
Dog               ............................................................
Fox               ............................................................
Cat               MFRGNRQSAELQHTGEASRRPTPTAGLGSKTREGPRRLAERRLRRGESPPAEQPRPIGGL
Mouse             ............................................................
Rabbit            ............................................................
Lion              ............................................................
Tiger             ............................................................
Pig               ............................................................

Human             ............................................................
Bat [Rousettus    ............................................................
Dog               ............................................................
Fox               ............................................................
Cat               QQPLGDWRKRKKAGRRVQSFLPAGTFVFLHRVCGVAPRDPGRRPLDPRPSRGDPGARDRQ
Mouse             ............................................................
Rabbit            ............................................................
Lion              ............................................................
Tiger             ............................................................
Pig               ............................................................

                                 1        10        20        30        40
Human             ...............MSQWYELQQLDSKFLEQVHQLYDDSFPMEIRQYLAQWLEKQDWEH
Bat               ...............MSQWYELQQLDSKFLEQVHQLYDDSFPMEIRQYLAQWLEKQDWEH
Dog               ...............MSQWYELQQLDSKFLEQVHQLYDDCFPMEIRQYLAQWLEKQDWEH
Fox               ...............MSQWYELQQLDSKFLEQVHQLYDDCFPMEIRQYLAQWLEKQDWEH
Cat               HCCASAHCCPTQRSEMSQWYELQQLDSKFLEQVHQLYDDSFPMEIRQYLAQWLEKQDWEH
Mouse             ...............MSQWFELQQLDSKFLEQVHQLYDDSFPMEIRQYLAQWLEKQDWEH
Rabbit            ...............MSQWYELQQLDSKFLEQVHQLYDDSFPMEIRQYLAQWLENQDWEH
Lion              ...............MSQWYELQQLDSKFLEQVHQLYDDSFPMEIRQYLAQWLEKQDWEH
Tiger             ...............MSQWYELQQLDSKFLEQVHQLYDDSFPMEIRQYLAQWLEKQDWEH
Pig               ...............MSQWYELQQLDSKFLEQVHQLYDDSFPMEIRQYLAQWLENQDWEH

                      50        60        70        80        90       100
Human             AANDVSFATIRFHDLLSQLDDQYSRFSLENNFLLQHNIRKSKRNLQDNFQEDPIQMSMII
Bat               AANDVSFATIRFHDLLSQLDDQYSRFSLENNFLLQHNIRKSKRNLQDNFQEDPIQMSMII
Dog               AANDVSFATIRFHDLLSQLDDQYSRFSLENNFLLQHNIRKSKRNLQDNFQEDPIQMSMII
Fox               AANDVSFATIRFHDLLSQLDDQYSRFSLENNFLLQHNIRKSKRNLQDNFQEDPIQMSMII
Cat               AANDVSFATIRFHDLLSQLDDQYSRFSLENNFLLQHNIRKSKRNLQDNFQEDPIQMSMII
Mouse             AAYDVSFATIRFHDLLSQLDDQYSRFSLENNFLLQHNIRKSKRNLQDNFQEDPVQMSMII
Rabbit            AANDVSFATILFHDLLSQLDDQYSRFSLENNFLLQHNIRKSKRNLQDNFQEDPIQMSMII
Lion              AANDVSFATIRFHDLLSQLDDQYSRFSLENNFLLQHNIRKSKRNLQDNFQEDPIQMSMII
Tiger             AANDVSFATIRFHDLLSQLDDQYSRFSLENNFLLQHNIRKSKRNLQDNFQEDPIQMSMII
Pig               AANDVSFATIRFHDLLSQLDDQYSRFSLENNFLLQHNIRKSKRNLQDNFQEDPIQMSMII

                      110       120       130       140       150       160
Human             YSCLKEERKILENAQRFNQAQSGNIQSTVMLDKQKELDSKVRNVKDKVMCIEHEIKSLED
Bat               YNCLKEERKILENAQRFNQAQSGNIQSTVMLDKQKELDSKVRNVKDKVMCIEHEIKTLED
Dog               YNCLKEERKILENAQRFNQAQSGNVQSTVMLDKQKELDSKVRNVKDKVMCIEHEIKTLED
Fox               YNCLKEERKILENAQRFNQAQSGNVQSTVMLDKQKELDSKVRNVKDKVMCIEHEIKTLED
Cat               YNCLKEERKILENAQRFNQAQSGNIQSTVMLDKQKELDSKVRNVKDKVMCIEHEIKTLED
Mouse             YNCLKEERKILENAQRFNQAQEGNIQNTVMLDKQKELDSKVRNVKDQVMCIEQEIKTLEE
Rabbit            YNCLKEERKILENAQRFNQAQSGNIQSTVMLDKQKELDNKVRNVKEQVMNIEHEIKTLED
Lion              YNCLKEERKILENAQRFNQAQSGNIQSTVMLDKQKELDSKVRNVKDKVMCIEHEIKTLED
Tiger             YNCLKEERKILENAQRFNQAQSGNIQSTVMLDKQKELDSKVRNVKDKVMCIEHEIKTLED
Pig               CNCLKEERKILENAQRFNQTQSGNIQSTVMLDKHKELDSKVRNVKDKVMCIEHEIKTLED

                      170       180       190       200       210       220
Human             LQDEYDFKCKTLQNREHETNGVAKSDQKQEQLLLKKMYLMLDNKRKEVVHKIIELLNVTE
Bat               LQDEYDFKCKTSQNREHEANGVAKNDQKQEQMLLHKMYLMLDNKRKEVVLKIIELLNITE
Dog               LQDEYDFKCKTLQNREHEANGVAKNDQKQEQMLIQKMYLMLDNKRKEVVLKIIELLNVTE
Fox               LQDEYDFKCKTLQNREHEANGVAKNDQKQEQMLIQKMYLMLDNKRKEVVLKIIELLNVTE
Cat               LQDEYDFKCKTLQNREHETNGVAKNDQKQEQMLIKKMYLMLDSKRKEVVHKIIELLNITE
Mouse             LQDEYDFKCKTSQNREGEANGVAKSDQKQEQLLLHKMFLMLDNKRKEIIHKIRELLNSIE
Rabbit            LQDEYDFKCKTLQNREHEANGVAKSDQKQEQLLLQKMYLMLDNKRKEIVHKMRELLNATE
Lion              LQDEYDFKCKTLQNREHETNGVAKNDQKQEQMLIKKMYLMLDSKRKEVVHKIIELLNITE
Tiger             LQDEYDFKCKTLQNREHETNGVAKNDQKQEQMLIKKMYLMLDSKRKEVVHKIIELLNITE
Pig               LQDEYDFKCKTLQNREHDTNGVAKNDQKQEQMLLQKMYLMLDNKRKEVVHKIIELLNVTE
```

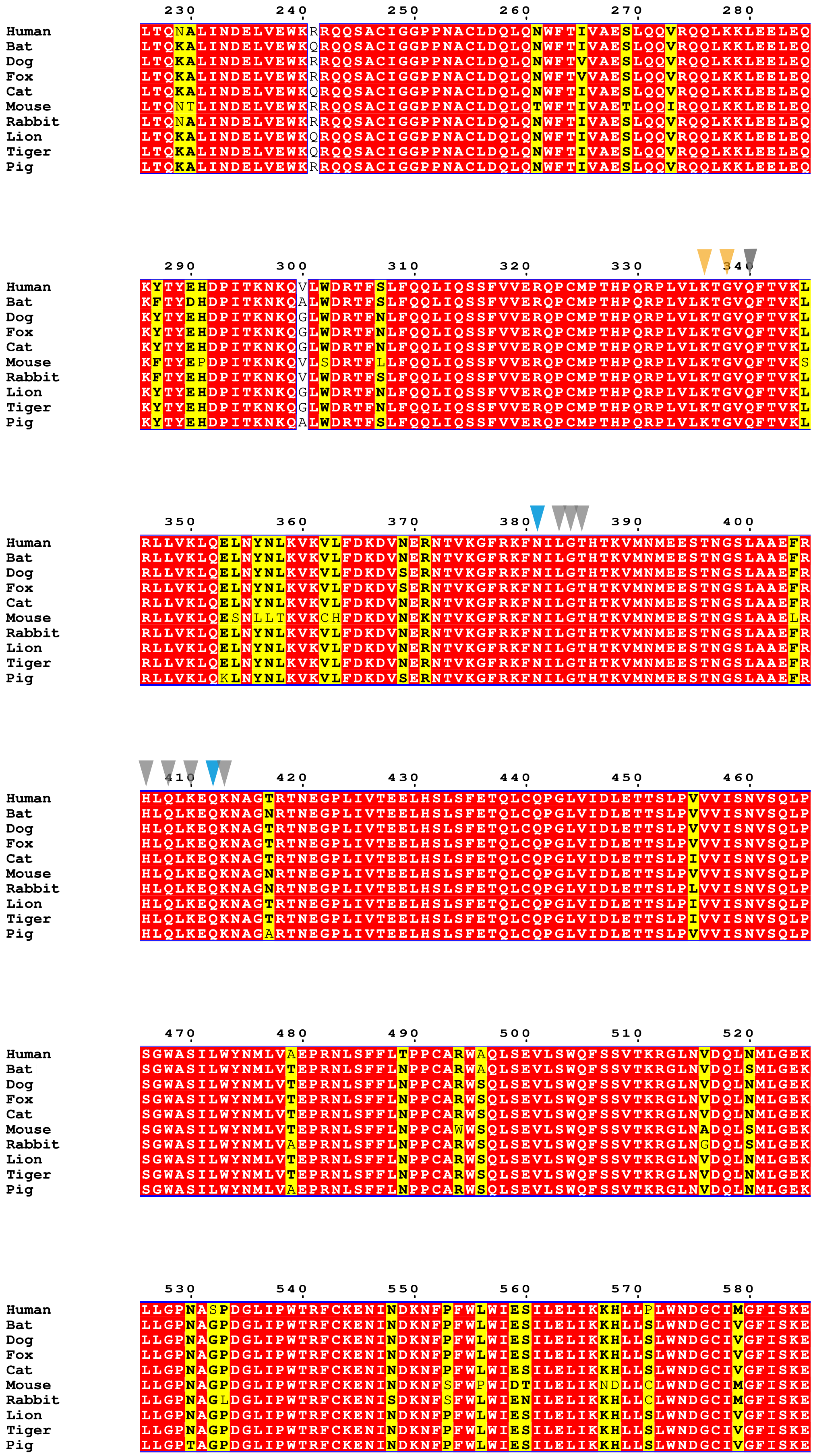
230 240 250 260 270 280
Human LTQNALINDELVEWKRRQQSACIGGPPNACLDQLQNWFTIVAESLQQVRQQLKKLEELEQ
Bat LTQKALINDELVEWKQRQQSACIGGPPNACLDQLQNWFTIVAESLQQVRQQLKKLEELEQ
Dog LTQKALINDELVEWKRRQQSACIGGPPNACLDQLQNWFTVVAESLQQVRQQLKKLEELEQ
Fox LTQKALINDELVEWKRRQQSACIGGPPNACLDQLQNWFTIVAESLQQVRQQLKKLEELEQ
Cat LTQKALINDELVEWKQRQQSACIGGPPNACLDQLQNWFTIVAESLQQVRQQLKKLEELEQ
Mouse LTQNTLINDELVEWKRRQQSACIGGPPNACLDQLQTWFTIVAETLQQIRQQLKKLEELEQ
Rabbit LTQNALINDELVEWKRRQQSACIGGPPNACLDQLQNWFTIVAESLQQVRQQLKKLEELEQ
Lion LTQKALINDELVEWKQRQQSACIGGPPNACLDQLQNWFTIVAESLQQVRQQLKKLEELEQ
Tiger LTQKALINDELVEWKQRQQSACIGGPPNACLDQLQNWFTIVAESLQQVRQQLKKLEELEQ
Pig LTQKALINDELVEWKRRQQSACIGGPPNACLDQLQNWFTIVAESLQQVRQQLKKLEELEQ
290 300 310 320 330 340
Human KYTYEHDPITKNKQVLWDRTFSLFQQLIQSSFVVERQPCMPTHPQRPLVLKTGVQFTVKL
Bat KFTYDHDPITKNKQALWDRTFSLFQQLIQSSFVVERQPCMPTHPQRPLVLKTGVQFTVKL
Dog KYTYEHDPITKNKQGLWDRTFNLFQQLIQSSFVVERQPCMPTHPQRPLVLKTGVQFTVKL
Fox KYTYEHDPITKNKQGLWDRTFNLFQQLIQSSFVVERQPCMPTHPQRPLVLKTGVQFTVKL
Cat KYTYEHDPITKNKQGLWDRTFNLFQQLIQSSFVVERQPCMPTHPQRPLVLKTGVQFTVKL
Mouse KFTYEPDPITKNKQVLSDRTFLLFQQLIQSSFVVERQPCMPTHPQRPLVLKTGVQFTVKS
Rabbit KFTYEHDPITKNKQVLWDRTFSLFQQLIQSSFVVERQPCMPTHPQRPLVLKTGVQFTVKL
Lion KYTYEHDPITKNKQGLWDRTFNLFQQLIQSSFVVERQPCMPTHPQRPLVLKTGVQFTVKL
Tiger KYTYEHDPITKNKQGLWDRTFNLFQQLIQSSFVVERQPCMPTHPQRPLVLKTGVQFTVKL
Pig KYTYEHDPITKNKQALWDRTFSLFQQLIQSSFVVERQPCMPTHPQRPLVLKTGVQFTVKL
350 360 370 380 390 400
Human RLLVKLQELNYNLKVKVLFDKDVNERNTVKGFRKFNILGTHTKVMNMEESTNGSLAAEFR
Bat RLLVKLQELNYNLKVKVLFDKDVNERNTVKGFRKFNILGTHTKVMNMEESTNGSLAAEFR
Dog RLLVKLQELNYNLKVKVLFDKDVSERNTVKGFRKFNILGTHTKVMNMEESTNGSLAAEFR
Fox RLLVKLQELNYNLKVKVLFDKDVSERNTVKGFRKFNILGTHTKVMNMEESTNGSLAAEFR
Cat RLLVKLQELNYNLKVKVLFDKDVNERNTVKGFRKFNILGTHTKVMNMEESTNGSLAAEFR
Mouse RLLVKLQESNLLTKVKCHFDKDVNEKNTVKGFRKFNILGTHTKVMNMEESTNGSLAAELR
Rabbit RLLVKLQELNYNLKVKVLFDKDVNERNTVKGFRKFNILGTHTKVMNMEESTNGSLAAEFR
Lion RLLVKLQELNYNLKVKVLFDKDVNERNTVKGFRKFNILGTHTKVMNMEESTNGSLAAEFR
Tiger RLLVKLQELNYNLKVKVLFDKDVNERNTVKGFRKFNILGTHTKVMNMEESTNGSLAAEFR
Pig RLLVKLQKLNYNLKVKVLFDKDVSERNTVKGFRKFNILGTHTKVMNMEESTNGSLAAEFR
410 420 430 440 450 460
Human HLQLKEQKNAGTRTNEGPLIVTEELHSLSFETQLCQPGLVIDLETTSLPVVVISNVSQLP
Bat HLQLKEQKNAGNRTNEGPLIVTEELHSLSFETQLCQPGLVIDLETTSLPVVVISNVSQLP
Dog HLQLKEQKNAGTRTNEGPLIVTEELHSLSFETQLCQPGLVIDLETTSLPVVVISNVSQLP
Fox HLQLKEQKNAGTRTNEGPLIVTEELHSLSFETQLCQPGLVIDLETTSLPVVVISNVSQLP
Cat HLQLKEQKNAGTRTNEGPLIVTEELHSLSFETQLCQPGLVIDLETTSLPIVVISNVSQLP
Mouse HLQLKEQKNAGNRTNEGPLIVTEELHSLSFETQLCQPGLVIDLETTSLPVVVISNVSQLP
Rabbit HLQLKEQKNAGNRTNEGPLIVTEELHSLSFETQLCQPGLVIDLETTSLPLVVISNVSQLP
Lion HLQLKEQKNAGTRTNEGPLIVTEELHSLSFETQLCQPGLVIDLETTSLPIVVISNVSQLP
Tiger HLQLKEQKNAGTRTNEGPLIVTEELHSLSFETQLCQPGLVIDLETTSLPIVVISNVSQLP
Pig HLQLKEQKNAGARTNEGPLIVTEELHSLSFETQLCQPGLVIDLETTSLPVVVISNVSQLP
470 480 490 500 510 520
Human SGWASILWYNMLVAEPRNLSFFLTPPCARWAQLSEVLSWQFSSVTKRGLNVDQLNMLGEK
Bat SGWASILWYNMLVTEPRNLSFFLNPPCARWAQLSEVLSWQFSSVTKRGLNVDQLSMLGEK
Dog SGWASILWYNMLVTEPRNLSFFLNPPCARWSQLSEVLSWQFSSVTKRGLNVDQLNMLGEK
Fox SGWASILWYNMLVTEPRNLSFFLNPPCARWSQLSEVLSWQFSSVTKRGLNVDQLNMLGEK
Cat SGWASILWYNMLVTEPRNLSFFLNPPCARWSQLSEVLSWQFSSVTKRGLNVDQLNMLGEK
Mouse SGWASILWYNMLVTEPRNLSFFLNPPCAWWSQLSEVLSWQFSSVTKRGLNADQLSMLGEK
Rabbit SGWASILWYNMLVAEPRNLSFFLNPPCARWSQLSEVLSWQFSSVTKRGLNGDQLSMLGEK
Lion SGWASILWYNMLVTEPRNLSFFLNPPCARWSQLSEVLSWQFSSVTKRGLNVDQLNMLGEK
Tiger SGWASILWYNMLVTEPRNLSFFLNPPCARWSQLSEVLSWQFSSVTKRGLNVDQLNMLGEK
Pig SGWASILWYNMLVAEPRNLSFFLNPPCARWSQLSEVLSWQFSSVTKRGLNVDQLNMLGEK
530 540 550 560 570 580
Human LLGPNASPDGLIPWTRFCKENINDKNFPFWLWIESILELIKKHLLPLWNDGCIMGFISKE
Bat LLGPNAGPDGLIPWTRFCKENINDKNFPFWLWIESILELIKKHLLSLWNDGCIVGFISKE
Dog LLGPNAGPDGLIPWTRFCKENINDKNFPFWLWIESILELIKKHLLSLWNDGCIVGFISKE
Fox LLGPNAGPDGLIPWTRFCKENINDKNFPFWLWIESILELIKKHLLSLWNDGCIVGFISKE
Cat LLGPNAGPDGLIPWTRFCKENINDKNFPFWLWIESILELIKKHLLSLWNDGCIVGFISKE
Mouse LLGPNAGPDGLIPWTRFCKENINDKNFSFWPWIDTILELIKNDLLCLWNDGCIMGFISKE
Rabbit LLGPNAGLDGLIPWTRFCKENISDKNFSFWLWIENILELIKKHLLCLWNDGCIMGFISKE
Lion LLGPNAGPDGLIPWTRFCKENINDKNFPFWLWIESILELIKKHLLSLWNDGCIVGFISKE
Tiger LLGPNAGPDGLIPWTRFCKENINDKNFPFWLWIESILELIKKHLLSLWNDGCIVGFISKE
Pig LLGPTAGPDGLIPWTRFCKENINDKNFPFWLWIESILELIKKHLLSLWNDGCIVGFISKE

```
                 590       600       610       620       630       640
Human    RERALLKDQQPGTFLLRFSESSREGAITFTWVERSQNGGEPDFHAVEPYTKKELSAVTFP
Bat      RERALLKDQQPGTFLLRFSESCREGAITFTWVERSQNGGEPYFHAVEPYTKKELSAVTFP
Dog      RERALLKDQQPGTFLLRFSESCREGAITFTWVERAQGGGEPYFHAVEPYTKKELSAVTFP
Fox      RERALLKDQQPGTFLLRFSESCREGAITFTWVERAQGGGEPYFHAVEPYTKKELSAVTFP
Cat      RERALLKDQQPGTFLLRFSESCREGAITFTWVERSQNGGEPYFHAVEPYTKKELSAVTFP
Mouse    RERALLKDQQPGTFLLRFSESSREGAITFTWVERSQNGGEPDFHAVEPYTKKELSAVTFP
Rabbit   RERALLKDQQPGTFLLRFSESSREGAITFTWVERSQNGGEPYFHAVEPYTKKELSAVTFP
Lion     RERALLKDQQPGTFLLRFSESCREGAITFTWVERSQNGGEPYFHAVEPYTKKELSAVTFP
Tiger    RERALLKDQQPGTFLLRFSESCREGAITFTWVERSQNGGEPYFHAVEPYTKKELSAVTFP
Pig      RERALLKDQQPGTFLLRFSESCREGAITFTWVERSQNGGEPYFHAVEPYTKKELSAVTFP

                 650       660       670       680       690       700
Human    DIIRNYKVMAAENIPENPLKYLYPNIDKDHAFGKYYSRPKEAPEPMELDGPKGTGYIKTE
Bat      DIIRNYKVMAAENIPENPLKYLYPNIDKDHAFGKYYSRPKEAPEPMELDGPKGTGYIKTE
Dog      DIIRNYKVMAAENIPENPLKFLYPNIDKDHAFGKYYSRPKEAPEPMELDGPKGTGYIKTE
Fox      DIIRNYKVMAAENIPENPLKFLYPNIDKDHAFGKYYSRPKEAPEPMELDGPKGTGYIKTE
Cat      DIIRNYKVMAAENIPENPLKYLYPNIDKDHAFGKYYSRPKEAPEPMELDGPKGTGYIKTE
Mouse    DIIRNYKVMAAENIPENPLKYLYPNIDKDHAFGKYYSRPKEAPEPMELDDPKRTGYIKTE
Rabbit   DIIRNYKVMAAENIPENPLKYLYPNIDKDHAFGKYYSRPKEGQEPMELDGPKGTGYIKTE
Lion     DIIRNYKVMAAENIPENPLKYLYPNIDKDHAFGKYYSRPKEAPEPMELDGPKGTGYIKTE
Tiger    DIIRNYKVMAAENIPENPLKYLYPNIDKDHAFGKYYSRPKEAPEPMELDGPKGTGYIKTE
Pig      DIIRNYKVMAAENIPENPLKYLYPNIDKDHAFGKYYSRPKEAPEPMELDGPKGTGYIKTE

                 710       720       730       740       750
Human    LISVSEVHPSRLQTTDNLLPMSPEEFDEVSRIVGSVEFDSMMNTV.......
Bat      LISVSEVHPSLLQTTDNLLPMSPEEFDEVSRIVGTVEFDSMMNV........
Dog      LISVSEVHPSRLQTTDNLLPMSPEEFDEVSRIVGSVEFDSMMHAV.......
Fox      LISVSEVHPSRLQTTDNLLPMSPEEFDEVSRIVGSVEFDSMMHAV.......
Cat      LISVSEVHPSRLQTTDNLLPMSPEEFDEVSRIVGPVEFDTMMNAV.......
Mouse    LISVSEVHPSRLQTTDNLLPMSPEEFDEMSRIVGPE.FDSMMSTV.......
Rabbit   LISVSEVHPSRLQSTENLLPMSPEEFDEVSRIMGPVEFDSMMNAV.......
Lion     LISVSEVHPSRLQTTDNLLPMSPEEFDEVSRIVGPVEFDTMMNAV.......
Tiger    LISVSEVHPSRLQTTDNLLPMSPEEFDEVSRIVGPVEFDTMMNAV.......
Pig      LISVSEVHPSRLQTTDNLLPMSPEEFDEVSRMVGPVEFDVTWNKFSGTMNLD
```

### Supplementary Figure S7

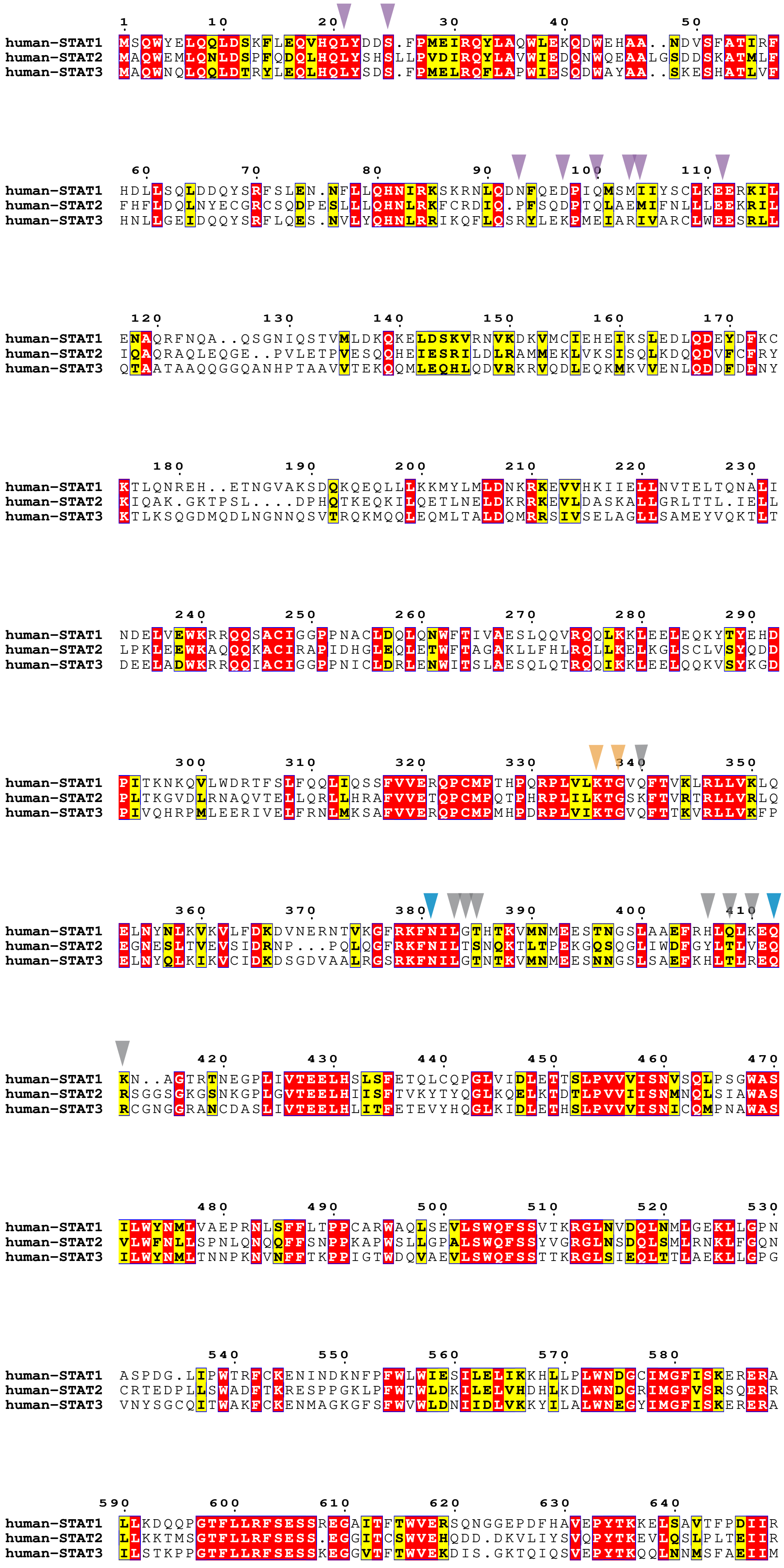

```
               650       660       670       680       690       700
human-STAT1    NYKVMAAENIPENPLKYLYPNIDKDHAFGKYYSRPKEAPEPMELDGPKGTGYIKTELISV
human-STAT2    HYQLLTEENIPENPLRFLYPRIPRDEAFGCYYQEKVNLQ........ERRKYLKHRLIVV
human-STAT3    GYKIMDATNILVSPLVYLYPDIPKEEAFGKYCRPES.QEHP.EADPGSAAPYLKTKFICV

               710       720                      730       740
human-STAT1    SEVHPSRLQTTDNLLPM............SPE..EFDEVSRIVGS.........VEFDSM
human-STAT2    SNRQVDELQQPLELKPEPELESLELELGLVPEPELSLDLEPLLKAGLDLGPELESVLEST
human-STAT3    TPTTCS....NTIDLPM............SPR..TLDSLMQFGNNGEGAEPSAGGQFESL

                  750
human-STAT1    MNTV........................................................
human-STAT2    LEPVIEPTLCMVSQTVPEPDQGPVSQPVPEPDLPCDLRHLNTEPMEIFRNCVKIEEIMPN
human-STAT3    TFDMELTSECATSPM.............................................

human-STAT1    .................................
human-STAT2    GDPLLAGQNTVDEVYVSRPSHFYTDGPLMPSDF
human-STAT3    .................................
```

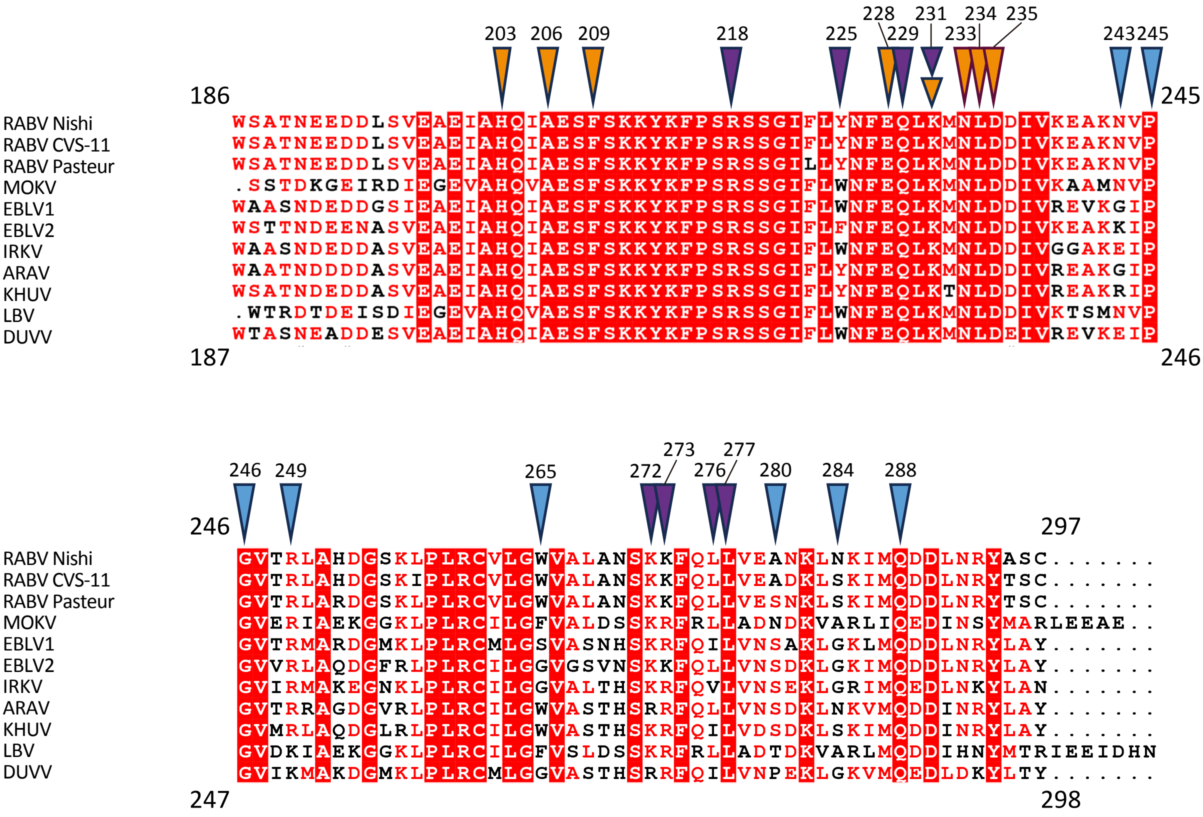
203 206 209 218 225 228 229 231 233 234 235 243 245
186
245
RABV Nishi WSATNEEDDLSVEAEIAHQIAESFSKKYKFPSRSSGIFLYNFEQLKMNLDDIVKEAKNVP
RABV CVS-11 WSATNEEDDLSVEAEIAHQIAESFSKKYKFPSRSSGIFLYNFEQLKMNLDDIVKEAKNVP
RABV Pasteur WSATNEEDDLSVEAEIAHQIAESFSKKYKFPSRSSGILLYNFEQLKMNLDDIVKEAKNVP
MOKV .SSTDKGEIRDIEGEVAHQVAESFSKKYKFPSRSSGIFLWNFEQLKMNLDDIVKAAMNVP
EBLV1 WAASNDEDDGSIEAEIAHQIAESFSKKYKFPSRSSGIFLWNFEQLKMNLDDIVREVKGIP
EBLV2 WSTTNDEENASVEAEIAHQIAESFSKKYKFPSRSSGIFLFNFEQLKMNLDDIVKEAKKIP
IRKV WAASNDEDDASVEAEIAHQIAESFSKKYKFPSRSSGIFLWNFEQLKMNLDDIVGGAKEIP
ARAV WAATNDDDDASVEAEIAHQIAESFSKKYKFPSRSSGIFLYNFEQLKMNLDDIVREAKGIP
KHUV WSATNDEDDASVEAEIAHQIAESFSKKYKFPSRSSGIFLYNFEQLKTNLDDIVREAKRIP
LBV .WTRDTDEISDIEGEVAHQVAESFSKKYKFPSRSSGIFLWNFEQLKMNLDDIVKTSMNVP
DUVV WTASNEADDESVEAEIAHQIAESFSKKYKFPSRSSGIFLWNFEQLKMNLDEIVREVKEIP
187
246
246 249 265 272 273 276 277 280 284 288
246
297
RABV Nishi GVTRLAHDGSKLPLRCVLGWVALANSKKFQLLVEANKLNKIMQDDLNRYASC.......
RABV CVS-11 GVTRLAHDGSKIPLRCVLGWVALANSKKFQLLVEADKLSKIMQDDLNRYTSC.......
RABV Pasteur GVTRLARDGSKLPLRCVLGWVALANSKKFQLLVESNKLSKIMQDDLNRYTSC.......
MOKV GVERIAEKGGKLPLRCILGFVALDSSKRFRLLADNDKVARLIQEDINSYMARLEEAE..
EBLV1 GVTRMARDGMKLPLRCMLGSVASNHSKRFQILVNSAKLGKLMQDDLNRYLAY.......
EBLV2 GVVRLAQDGFRLPLRCILGGVGSVNSKKFQLLVNSDKLGKIMQDDLNRYLAY.......
IRKV GVIRMAKEGNKLPLRCILGGVALTHSKRFQVLVNSEKLGRIMQEDLNKYLAN.......
ARAV GVTRRAGDGVRLPLRCILGWVASTHSKRFQLLVNSDKLNKVMQDDINRYLAY.......
KHUV GVMRLAQDGLRLPLRCILGWVASTHSKRFQILVDSDKLSKIMQDDINRYLAY.......
LBV GVDKIAEKGGKLPLRCILGFVSLDSSKRFRLLADTDKVARLMQDDIHNYMTRIEEIDHN
DUVV GVIKMAKDGMKLPLRCMLGGVASTHSRRFQILVNPEKLGKVMQEDLDKYLTY.......
247
298

**Supplementary Figure S1 Single-particle cryo-EM analysis workflow of pY-STAT1/P-CTD complex.**

Representative cryo-EM micrograph and 2D class images (left). Cryo-EM data processing flowchart for pY-STAT1-P-CTD complex (right). Focused masks used for 3D classification for the tetramer reconstruction are presented in transparent brown.

**Supplementary Figure S2 Cryo-EM reconstruction of pY-STAT1/ P-CTD complex**

**(A)** Angular distribution of the particles used in 3D reconstruction (pY-STAT1/P-CTD complex (pY-STAT1 dimer)). **(B)** FSC curve of the density map for the pY-STAT1/P-CTD complex (pY-STAT1 dimer) calculated by relion_postprocess[4] .

**(C)** Local resolution map of the pY-STAT1/P-CTD complex (pY-STAT1 dimer). **(D)** Orientation distribution map of the pY-STAT1/P-CTD complex (pY-STAT1 /P-CTD complex (pY-STAT1 tetramer)). **(E)** FSC curve of the density map for the pY-STAT1/P-CTD complex (pY-STAT1 tetramer). **(F)** Local resolution map of the pY-STAT1/P-CTD complex (pY-STAT1 tetramer). **(B, E)** The resolution at FSC = 0.143 is indicated. **(C, F)** The color bar indicates local resolution in Å.

**Supplementary Figure S3 Sample preparation and Cryo-EM analysis of pY-STAT1–P-CTD complex**

**(A)** SEC chromatogram of pY-STAT1−P-CTD complex. **(B)** SDS-PAGE analysis confirming the formation and purity of the pY-STAT1 − P-CTD complex fractionated by SEC chromatography shown (in (**A**)). Molecular weight of each marker proteins is indicated. **(C)** 2D class averages of the pY-STAT1/P-CTD tetramer. Red arrows indicate locations of NTD dimers. **(D)** EM density around phosphorylated Tyr701 of protomer B (shown in dark green with phosphate moiety of orange (phosphorous atom) and red (oxygen atoms)) recognized by the reciprocal SH2 domain of protomer A (light green). The side chains of Lys584, Arg602 of protomer B are labeled, together with the main chain amide nitrogen atoms of Glu605 and Ser606. Hydrogen bonds are shown as dashed lines.

**Supplementary Figure S4 Residues mediating the pY-STAT1/P-CTD interaction (A)–(J)** Amino acid residues involved in the interaction between pY-STAT1 and P-CTD. The interactions in the A site are shown in (A)–(C), those in the B site are in (D)–(F), and those in the C site are in (G) and (H). The EM map of each site is shown as mesh, and the model is colored as in Fig. 1d. Hydrogen bonds and salt bridges are indicated by yellow dashed lines. (**I**) The location of Asn226 of P-CTD. The side chains of Glu228 interacting with the pY-STAT1 A protomer, Gln229 interacting with the pY-STAT1 B protomer, and Ser210 that undergoes phosphorylation are also shown. pY-STAT1 A protomer is shown in light green, pY-STAT1 B protomer in magenta, and P-CTD in blue.

**Supplementary Figure S5 CD spectra comparison of the RABV P-CTD wild type and mutants**

**a.** Spectra of point mutants of either the A or B site. Each spectrum is shown as follows, P-CTD wild type, blue; F209A, red; W265G, purple. W265G is the same data as previously reported[5]. **b.** Spectra of mutants of the C site. Each spectrum is shown as follows, P-CTD wild type, blue; R218Q, gray; R218Q/Q229N/K231S/K273S, yellow.

**Supplementary Figure S6 Multiple sequence alignment of mammalian STAT1** Alignment (Clustal Omega) was performed for STAT1 of Human (*Homo sapiens*, uniprot, P42224), Bat (*Rousettus aegyptiacus*, C9K6U7), Dog (*Canis lupus familiaris*, A0A8I3PX00), Fox (*Vulpes vulpes*, A0A3Q7TB43), Cat (*Felis catus*, A0A337SVH3), Mouse (*Mus musculus*, P42225), Rabbit (*Oryctolagus cuniculus*, G1SHL8), Lion (*Panthera leo*, A0A8C8XVG1), Tiger (*Panthera tigris altaica*, A0A8C9JQ80), Pig (*Sus scrofa*, Q764M5). Residues conserved completely are highlighted in red with white letters; residues partially conserved are highlighted in yellow (conserved residues are shown in bold black letters). Residues of human STAT1 that interact with P-CTD are indicated with colored-triangles on top sequences (human STAT1) as follows. Residues used for the P-CTD interaction at both A site in A protomer and B site in B protomer are indicated in gray. Residues used for the interaction only at the A site are colored in orange. Residues used for the interaction only at the B site are colored in blue. Residues used for the interaction only at the C site are colored in purple.

**Supplementary Figure S7 Sequence alignment of human STAT1, STAT, and STAT3.** Human STAT1 (P42224), STAT2 (P52630), and STAT3 (P40763) are aligned (Clustal Omega). Amino acids conserved completely are highlighted in red with white letters; partially conserved amino acids are highlighted in yellow with bold black letters. Residues used for the interaction with P-CTD, indicated with triangles on the top sequence (STAT1) are indicated with the same coloring as in Fig. S6.

**Supplementary Figure S8 Multiple sequence alignment of the lyssavirus P-CTD.** Alignment (Clustal Omega) was performed for P-CTDs of RABV Nishi (Nishigahara strain, SWISSPROTdb Q9IPJ8), RABV CVS (CVS strain, P22363), RABV Pasteur (Pasteur vaccine strain (PV), P06747), MOKV (Mokola virus, P0C569), EBLV1 (European bat lyssavirus 1, A4UHP9), EBLV2 (European bat lyssavirus 2, A4UHQ4), IRKV (Irkut virus, Q5VKP5), ARAV (Aravan virus, Q6X1D7), KHUV (Khujand virus, Q6X1D3), MOKV (Mokola virus, P0C569), LBV (Lagos bat virus, O56773), and DUVV (Duvenhage virus, O56774). Amino acids conserved completely are highlighted in red with white letters; partially conserved residues are indicated as red letters. Residues used for the interaction with pY-STAT1 at the A site are shown with orange triangles, at the B site with blue triangles, and at the C site with purple triangles.
