## Supplemental Table for "Rabies virus antagonizes interferon signaling by targeting phosphorylated STAT1 tetramers"

**Supplementary table 1. Cryo-EM data collection, refinement and validation statistics**

| | pY-STAT1 dimer<br>(EMDB-82067)<br>(PDB 43QG) | pY-STAT1 tetramer<br>(EMDB-82102)<br>(PDB 43RL) |
|---|---|---|
| **Data collection and processing** | | |
| Magnification | 130 K | |
| Voltage (kV) | 300 | |
| Electron exposure (e–/Å$^2$) | 52.0 (dataset 1)<br>56.9 (dataset 2)<br>52.1 (dataset 3) | |
| Defocus range (μm) | -0.8 to -1.8 (dataset 1)<br>-0.8 to -1.4 (dataset 2)<br>-0.8 to -1.8 (dataset 3) | |
| Pixel size (Å) | 0.67 | |
| Symmetry imposed | *C*1 | *C*1 |
| Initial particle images (no.) | 581,276 (dataset 1)<br>2,943,912 (dataset 2)<br>722,719 (dataset 3) | |
| Final particle images (no.) | 97,930 | 27,081 |
| Map resolution (Å) | 3.1 | 3.7 |
| FSC threshold | | |
| **Refinement** | | |
| Initial model used (PDB code) | 7C20, 8YYV | 7C20, 8YYV |
| Model resolution (Å) | | |
| FSC threshold | 3.1 | 3.75 |
| Model composition | | |
| Non-hydrogen atoms | 11,006 | 21,861 |
| Protein residues | 1,351 | 2,660 |
| *B* factors (Å$^2$) | | |
| Protein | 31.05 | 65.71 |
| R.m.s. deviations | | |
| Bond lengths (Å) | 0.003 | 0.003 |
| Bond angles (°) | 0.591 | 0.657 |
| Validation by MolProbity | | |
| MolProbity score | 2.13 | 2.34 |
| Clashscore | 5.83 | 8.77 |
| Rotamer outliers (%) | 3.10 | 3.84 |
| Ramachandran plot | | |
| Favored (%) | 93.28 | 93.49 |
| Allowed (%) | 5.66 | 5.81 |
| Outliers (%) | 1.06 | 0.70 |
